# pH-driven evolutionary divergence and ongoing speciation in the cosmopolitan cyanobacterium *Microcoleus vaginatus*

**DOI:** 10.1101/2025.08.22.671682

**Authors:** Jingyi Wei, Hua Li, Haijian Yang, Renhui Li, Wensheng Shu, Chunxiang Hu

**Author notes:** Corresponding Authors: Hua Li and Chunxiang Hu.; Address: South Donghu Road 7, 430072 Wuhan, China. **Competing interests:** Authors declare that they have no competing interests.

## Abstract

Speciation models and the concept of speciation continuum have been well established in multicellular eukaryotes. However, the mechanisms underlying population differentiation and their environmental drivers in prokaryotes remain elusive. Here, we explored the population diversification of the filamentous cyanobacterium *Microcoleus vaginatus*, a photoautotrophic edificator with significant importance for dryland ecosystems worldwide. By constructing a 132-genome dataset spanning multiple climate zones across four continents, we integrated intraspecific genomics with environmental association analyses to elucidate the population genetic structure and molecular mechanisms underlying the divergence of the *M. vaginatus* continuum. Our findings reveal that adaptive differentiation is driven by the combined effects of natural selection, homologous recombination, and gene family turnover, with varying contributions across different phylogroups. The evolution of core and accessory genomes exhibits complementary patterns, with genome-wide association studies demonstrating the predominant role of the core genome in population divergence. Selection, followed by neutral evolution and recombination, mediates trade-offs in life-history strategies, while environmental factors shape these evolutionary processes. Notably, while local edaphic variables exert direct influences on Tajima’s D values, they demonstrate predominantly indirect modulation through alterations in the selection-to-gene gain ratio (S/G). pH emerges as the principal abiotic driver governing population genetic signatures, operating through coordinated effects on both S/G and the proportion of selected genes undergoing recombination (SUR). This study offers novel insights into the diversification mechanisms of cosmopolitan, dominant cyanobacteria, thereby advancing our understanding of prokaryotic speciation in complex natural settings and providing a framework for predicting microbial adaptation in a changing world.

## Introduction

Speciation has been a fundamental inquiry in evolutionary biology since the era of Darwin, with profound implications for understanding the origins and maintenance of biodiversity. The biological species concept, traditionally defined by reproductive isolation between phylogenetically proximate taxa, has been widely accepted for many decades. However, ongoing debates over the extent of reproductive isolation required have prompted the development of innovative conceptual models (Bobay and Ochman 2017a), including the speciation-with-gene-flow paradigm and the speciation continuum hypothesis (Wu 2001; Stankowski and Ravinet 2021). While these frameworks have gained empirical validation in multicellular eukaryotes (Papadopulos, et al. 2019; Wang, et al. 2024), investigations in prokaryotes remain notably underrepresented.

Since prokaryotes reproduce exclusively through asexual mechanisms, species delineation has historically relied upon 16S ribosomal RNA gene (*rrs*) similarity and whole-genome comparative thresholds. However, the pervasive occurrence of horizontal gene transfer (HGT) confers prokaryotes quasi-sexual characteristics, significantly reshaping evolutionary dynamics. In bacterial populations with large effective sizes, speciation emerges primarily through the intricate interplay between natural selection and recombination processes (Shapiro and Polz 2015; Shapiro 2018). The ubiquity of HGT, coupled with gene loss events that facilitate micro-niche differentiation and characterize bacterial lineages (Chu, et al. 2021), underscores the substantial contribution of accessory genomes to the speciation process. Nonetheless, researchers argue that functional constraints and selective pressures on the core genome constitute the predominant evolutionary drivers (Wang, et al. 2020). Lineage divergence typically emerges through the synergistic influence of environmental factors and geographic separation (Whittaker and Rynearson 2017; Li, et al. 2022). However, geographic barriers do not invariably prevent gene flow, thus generating reticulate evolutionary patterns among diverging lineages under sustained genetic exchange (Diop, et al. 2022; Dmitrijeva, et al. 2024). These findings highlight the multifaceted nature of evolutionary forces that govern species differentiation through diverse, environment-dependent mechanisms. Nevertheless, we still poorly understand how distinct evolutionary forces contribute to bacterial speciation and to what extent environmental factors modulate these processes.

*Microcoleus vaginatus* Gomont represents a bundle-forming filamentous cyanobacterium that exhibits dominance across dryland topsoils (approx. 41% of Earth’s land surface) worldwide while maintaining widespread distribution within aquatic microbial mats (Dvorak, et al. 2012; Hasler, et al. 2012). Distinguished by its remarkable sand-stabilizing capabilities and the biosynthesis of unique polysaccharides (Dembitsky, et al. 2001; Hu, et al. 2012), this species plays a crucial role in maintaining soil multifunctionality and facilitating the restoration of dryland ecosystems (Hu, et al. 2012; Rossi, et al. 2017). As the most abundant single-species terrestrial cyanobacterium documented to date (Moreira, et al. 2021), *M. vaginatus* is an ideal model for exploring ecological adaptation and evolutionary mechanisms in microbial populations. Traditional taxonomic approaches have long regarded *M. vaginatus* as a monotypic species based on *rrs* sequences and morphological traits. However, internal transcribed spacer sequence analyses have uncovered at least two phylogenetically distinct lineages adapted to moisture gradients (Hasler, et al. 2012). Contemporary genomic studies have revealed extensive genetic diversification and selective pressures within the *Microcoleus* genus, suggesting a global speciation continuum (Stanojkovic, et al. 2024). Despite recognizing distinct ecotypes within *M. vaginatus*, the mechanisms underlying speciation and evolutionary dynamics across diverse climatic regimes are still uncertain.

To address this knowledge gap, we sampled biological soil crusts (i.e., biocrusts) along a 3,400 km transect from temperate deserts and desert-steppe climatic zones throughout northwestern China, complemented by collections of cyanobacterial mats from subtropical humid and plateau mountain environments in southern China and the Tibetan Plateau. We employed an intraspecies whole-genome sequencing of *M. vaginatus* isolated from field collections and strains deposited in the Freshwater Algae Culture Collection at the Institute of Hydrobiology, Chinese Academy of Sciences (FACHB, https://algae.ihb.ac.cn/). By thoroughly integrating publicly available genomes from diverse biogeographic regions, encompassing European temperate, oceanic, and continental zones, Mediterranean climates, African tropical rainforests, oceanic ecosystems, and North American temperate, continental, and subtropical humid regions, we assembled a global genomic repository comprising 132 high-quality *M. vaginatus* intraspecific genomes.

By using population genomics methods coupled with comprehensive macro- and micro-environmental association analyses, we sought to illuminate (**1**) the phylogenetic architecture and genetic underpinnings of *M. vaginatus* population divergence; (**2**) the mechanistic interaction whereby natural selection, homologous recombination, and gene family turnover orchestrate adaptive divergence under ongoing gene flow; and (**3**) the environmental regulatory frameworks governing genomic evolutionary trajectories. We hypothesize that the ecological ascendancy of the nascent *M. vaginatus* across global biocrusts and aquatic mats reflects sophisticated adaptive evolutionary strategies analogous to habitat-mediated transitions in life-history paradigms (Dvorak, et al. 2012; Chen, et al. 2021; Li, et al. 2022). This divergence likely arises through selection pressures and functional gene reorganization in response to both edaphic and climatic gradients within the microhabitat and across geographic scales, potentially involving complementary evolutionary dynamics between core and accessory genomic components.

## Results and Discussion

### Genomic diversity reveals substantial intraspecific genetic plasticity

We constructed 58 high-quality *Microcoleus* genomes (completeness: 99.53 ± 0.42%; contamination: 0.2%; **Table S1**) from biocrusts and algal mats spanning 29° to 45°N and 87° to 115°E across China. The majority of genome assemblies were circularized into complete chromosomes (circularization rate > 75%), with N50 values exceeding 6 Mb. Through the incorporation of microscopic morphological observations, full-length *rrs* sequence alignment (at a 97% threshold), and an 11-bp conserved insert sequence (Garcia-Pichel, et al. 2001), we preliminarily classified these genomes as either *M. vaginatus* (57 strains) or *Microcoleus anatoxicus* (one strain). We then integrated 75 publicly available *M. vaginatus* genomes containing the diagnostic 11-bp conserved sequence, yielding a global dataset of 132 strains across four continents (**Fig. 1A**).

**Fig. 1.**
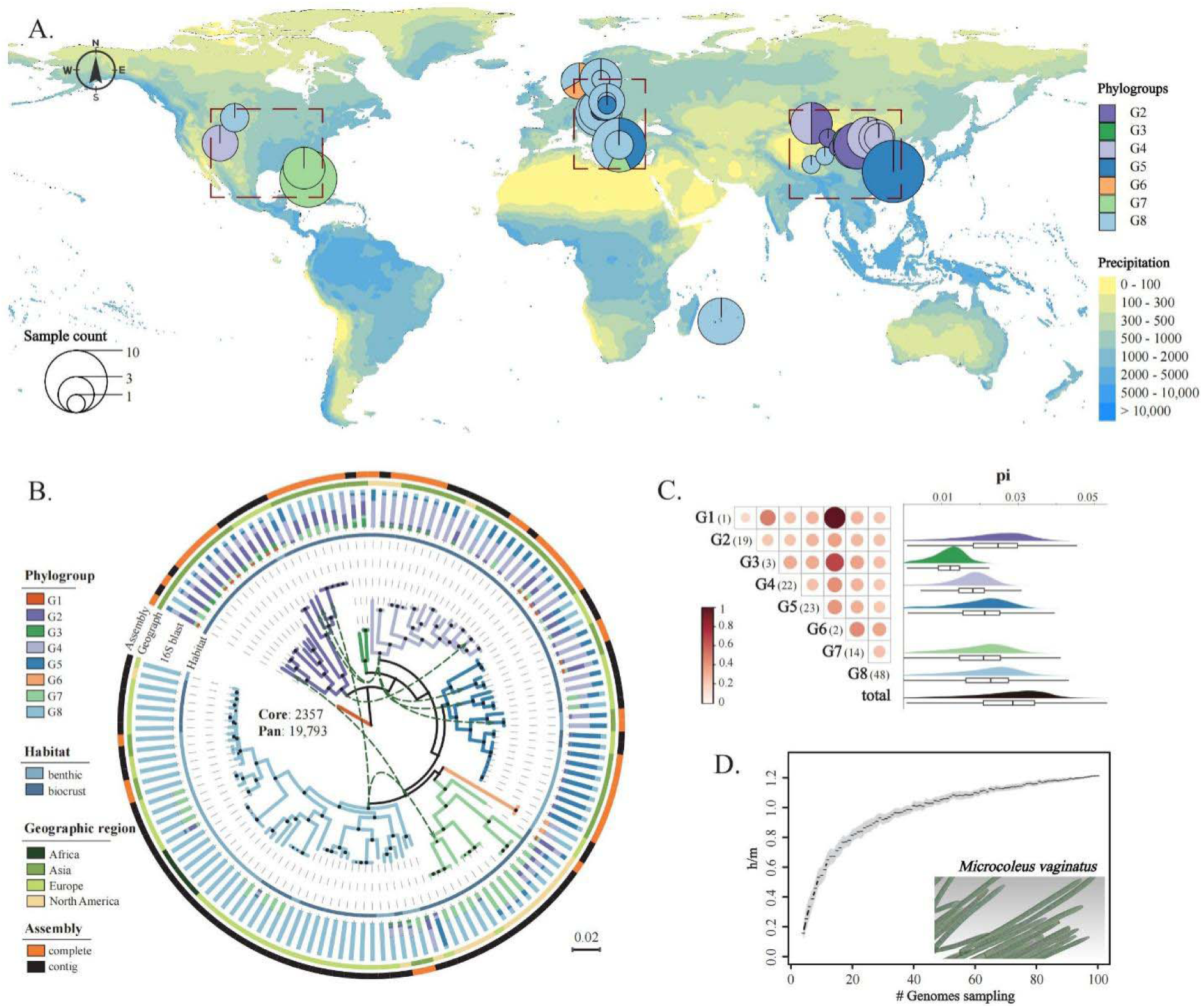
Phylogenetic architecture and intraspecific diversity patterns of *M. vaginatus*. (**A**) Global biogeographic distribution of sampled cyanobacterial strains across four continents. The color gradient of the map represents the annual precipitation, and the size of the circles corresponds to the number of samples collected at each site. The pie charts within the circles illustrate the relative abundance of different phylogenetic groups. (**B**) Maximum likelihood phylogenetic tree constructed from concatenated single-copy core protein sequences of 132 genomes, with distinct phylogroups delineated by differential branch coloration. Concentric rings from interior to exterior represent habitat classification, geographic origin, 16S rDNA alignment proportion, and genome assembly completeness, respectively. Solid black circles at nodes indicate bootstrap support values of 75% or higher, with the diameter of the circle proportional to the statistical confidence. Central numerical annotations denote core gene family counts and pan-genome family totals. Green dashed arrows highlight notable incongruencies between 16S rDNA sequence similarities and phylogenetic relationships. (**C**) Matrix visualization of fixation indices (Fst) between phylogroups and nucleotide diversity (π) within phylogroups, with color intensity gradient from white to red representing increasing genetic differentiation. (**D**) Trajectory of homoplastic to non-homoplastic allele ratios (h/m) corresponding to increasing genome sample sizes. Homoplastic alleles represent SNPs exhibiting phylogenetically independent distribution patterns scattered across the evolutionary tree, whereas non-homoplastic alleles predominantly cluster on closely related phylogenetic branches, reflecting vertical inheritance patterns.

Genomic characterization revealed that 30 strains harbored multiple ribosomal operon sets with substantial sequence divergence, generating a secondary peak at approximately 80% in the *rrs* similarity distribution (**Fig. S1**). Further investigation demonstrated that these strains corresponded to non-circularized, contig-level genomes sourced from public databases, with *rrs* sequences distributed across different contigs within each genome. The reduced rRNA gene similarity could result from inadequate quality control during genome assembly or the horizontal acquisition of exogenous *rrs* sequences (Acinas, et al. 2004). After removing outlier sequences, 53 strains (including 50 sequenced in this study) retained 2 to 5 multiple *rrs* copies, with similarity peaks of ∼99%, but their whole-genome average nucleotide identity (ANI) ranged from 90.3% to 100% (**Fig. S1**). The core genome comprised 2357 gene families, whereas the pan-genome extended to 19,793 families, indicating that *M. vaginatus* simultaneously exhibits high genetic conservation and diversity. These findings support the microbial pan-genome theory (McInerney, et al. 2017), which proposes that species maintain essential functions through a limited set of conserved core genes while achieving niche differentiation via the extensive diversification of both core and non-core genetic elements. Although multiple *rrn* operons are frequently associated with enhanced growth rate (Romillac and Santorufo 2021), we observed no significant correlation between *rrs* copy number and minimal doubling time (**Fig. S2**). It suggests that *M. vaginatus* allocates considerable resources toward stress tolerance mechanisms rather than exclusively optimizing growth rates (Li, et al. 2022). Therefore, *M. vaginatus* sustains genetic stability and ecological adaptation through diversified strategies, according to a “key function conservation-large genetic plasticity” framework (McInerney, et al. 2017).

### Population divergence accords with speciation models under gene flow

To elucidate the phylogenetic relationships within *M. vaginatus*, we constructed a phylogenetic tree using *M. anatoxicus* CHAB4116 as the outgroup (**Fig. S3**), which exhibited an average *rrs* similarity of 96.7% and an average ANI of 83.1% with *M. vaginatus* strains. Then, we constructed a re-rooted phylogeny using this reference tree, based on 761,569 aligned amino acid sites from core single-copy genes (**Fig. 1B**). The resulting phylogeny revealed eight distinct monophyletic phylogroups (G1-G8), consistent with the recently reported cryptic multi-species phenomenon within the *Microcoleus* complex (Stanojkovic, et al. 2024). To further validate the reliability of these monophyletic groups, we employed the fastBAPS clustering method (**Fig. S4**), which corroborated the tree topology, except for G2, as this clade was partitioned into two sub-clusters due to substantial internal diversity. In addition, we constructed a gene tree based on nucleotide sequences of core single-copy genes (**Fig. S5A**), where G1 through G8 maintained monophyletic integrity. However, topological discrepancies appeared in G6 and G7, primarily stemming from differential codon usage preferences among phylogroups (**Fig. S5B**). The PCA analysis revealed that PC1 predominantly loaded the histidine codon CAC, while PC2 primarily represented the stop codon TAA, suggesting that phylogroup separation may be intrinsically linked to variations in growth rate. The phylogenetic network further demonstrated frequent gene flow during early evolutionary stages, which decreased significantly in recent periods (**Fig. S6**). These multi-methodological validations proved that *M. vaginatus* comprises eight diverging monophyletic intraspecific lineages.

Although geographic regions and ecotypes did not correspond strictly with phylogroups, the strains of G1-G4 predominantly originated from Asian terrestrial habitats, whereas G5-G8 strains were primarily derived from Europe, North America, and Africa with a higher proportion of benthic habitats (**Fig. 1B**). The interspersed distribution of terrestrial and aquatic strains throughout the phylogeny suggests that environmental adaptations in *M. vaginatus* likely occur during recent, multiple independent evolutionary events. We further constructed a phylogeny based on *rrs* sequences (**Fig. S7**) and calculated the proportion of each phylogroup within the top 10% of *rrs* pairwise alignment scores for individual strains (**Fig. 1B**). The results revealed a mosaic clustering pattern, wherein strains frequently clustered with members from other phylogroups while exhibiting lower similarity to strains within their phylogroup. It reflects the asynchronous evolution between highly conserved *rrs* sequences and core protein sequences, indicating that evolutionary rates are decoupled between housekeeping genes and other essential genes (Lv, et al. 2015). Analysis of genetic differentiation indices revealed that the median fixation index (Fst) between phylogroups was 0.30 (ranging from 0.15 to 0.44; **Fig. 1C**), characteristic of high-level genetic differentiation (Wright 1949). Excluding G1, G3, and G6 due to limited sample sizes, intra-phylogroup nucleotide diversity (π) ranked as G2 > G8 > G5 > G7 > G4. Within 10-kb sliding windows, approximately 200 variant sites were observed, reaching up to 500 in highly variable regions. Consequently, *M. vaginatus* exhibits substantial intra-population divergence, with the highly heterogeneous G2 and G8 lineages likely undergoing active adaptive radiation in arid and semi-humid environments (Filatov, et al. 2021).

Notably, *rrs* similarities between phylogroups consistently exceeded 98.5%, while ANI values often fell below 95% even within individual phylogroups (**Fig. S8A**). This discrepancy generates conflicting species delineations across different levels (Konstantinidis and Tiedje 2005; Stackebrandt and Ebers 2006), leading many researchers to classify *M. vaginatus* as multiple distinct species (Skoupy, et al. 2024; Stanojkovic, et al. 2024). Given that genetic isolation does not constitute a simple function of genomic concordance (Bobay and Ochman 2017a), we calculated the ratio of homoplasic to non-homoplasic alleles (h/m) to assess the intensity of gene flow. The h/m ratio showed a gradual elevation with increasing genome sample sizes (**Fig. 1D**). However, the inclusion of an exogenous genome (i.e., *M. anatoxicus* CHAB4116) produced a pronounced mid-range discontinuity (**Fig. S8B**), indicating the absence of gene flow barriers within *M. vaginatus*. Gene flow fraction analysis (**Fig. S9A**) further demonstrated that despite substantial divergence, an average of 38.26% of the core genome continues to be exchanged between lineages (peaking at 62.09%), far exceeding the interspecies introgression level (Diop, et al. 2022). Early and recent core genomic exchanges averaged 25.47% and 12.78%, respectively, consistent with phylogenetic network patterns (**Fig. S6**). Regardless of temporal dynamics, G2 and G7 predominantly functioned as recipients of gene flow, while G4, G5, and G8 primarily served as donors. Similarly, HGT events between phylogroups ranged from 10 to 129 (**Fig. S9B**), generally exceeding the HGT frequencies observed between species within the *Microcoleus* genus, even when sharing the same hosts (Stott, et al. 2024). HGT occurred frequently among phylogroups G2, G5, G7, and G8 but was less prevalent between G4 and others. Although the feasibility of speciation under gene flow remains a debated issue (Wolf and Ellegren 2017), recent studies using high-throughput sequencing approaches have provided empirical support for this hypothesis (Wu 2001; Papadopulos, et al. 2019; Wang, et al. 2024). Therefore, population differentiation in *M. vaginatus* likely represents ongoing speciation processes that conform to models of gene flow-mediated evolution.

### Divergent evolutionary strategies for core and accessory genomic regions

We inferred critical evolutionary events across core and accessory genomic regions to dissect the population differentiation mechanisms underlying the evolution of *M. vaginatus*. Three phylogroups (G1, G3, and G6) were excluded from subsequent analyses due to insufficient sample size (≤ 3 individuals per group). We initially quantified the nonsynonymous to synonymous substitution ratio (Ka/Ks) for core single-copy genes to identify lineage-specific genes under positive natural selection (PSGs). Variable numbers of PSGs were detected across phylogroups (**Fig. 2A**), in which G2 (n = 46) and G8 (n = 38) exhibited the highest counts, followed by G4 (n = 25), G5 (n = 24), and G7 (n = 18), substantially exceeding the 19 PSGs previously reported in *Microcoleus* (Stanojkovic, et al. 2024). The proportion of positively selected sites relative to gene length showed no significant variation among phylogroups. The phylogroup G2, despite harboring the highest PSG count, displayed a relatively restricted geographic distribution. In contrast, the remaining four phylogroups followed the expected correlation between broader distribution and stronger positive selection. Although intensive selection pressure can significantly compromise the accuracy of phylogenetic reconstruction (Stott and Bobay 2020), tree topologies remained consistent regardless of whether PSG was included or excluded (**Fig. S10**). Extensive PSG overlaps were observed among all phylogroups, with over 84% occurring between G2 and other phylogroups. It meant the number of lineage-specific PSGs was broadly high, but convergent selection under different environmental pressures was also considerable.

**Fig. 2.**
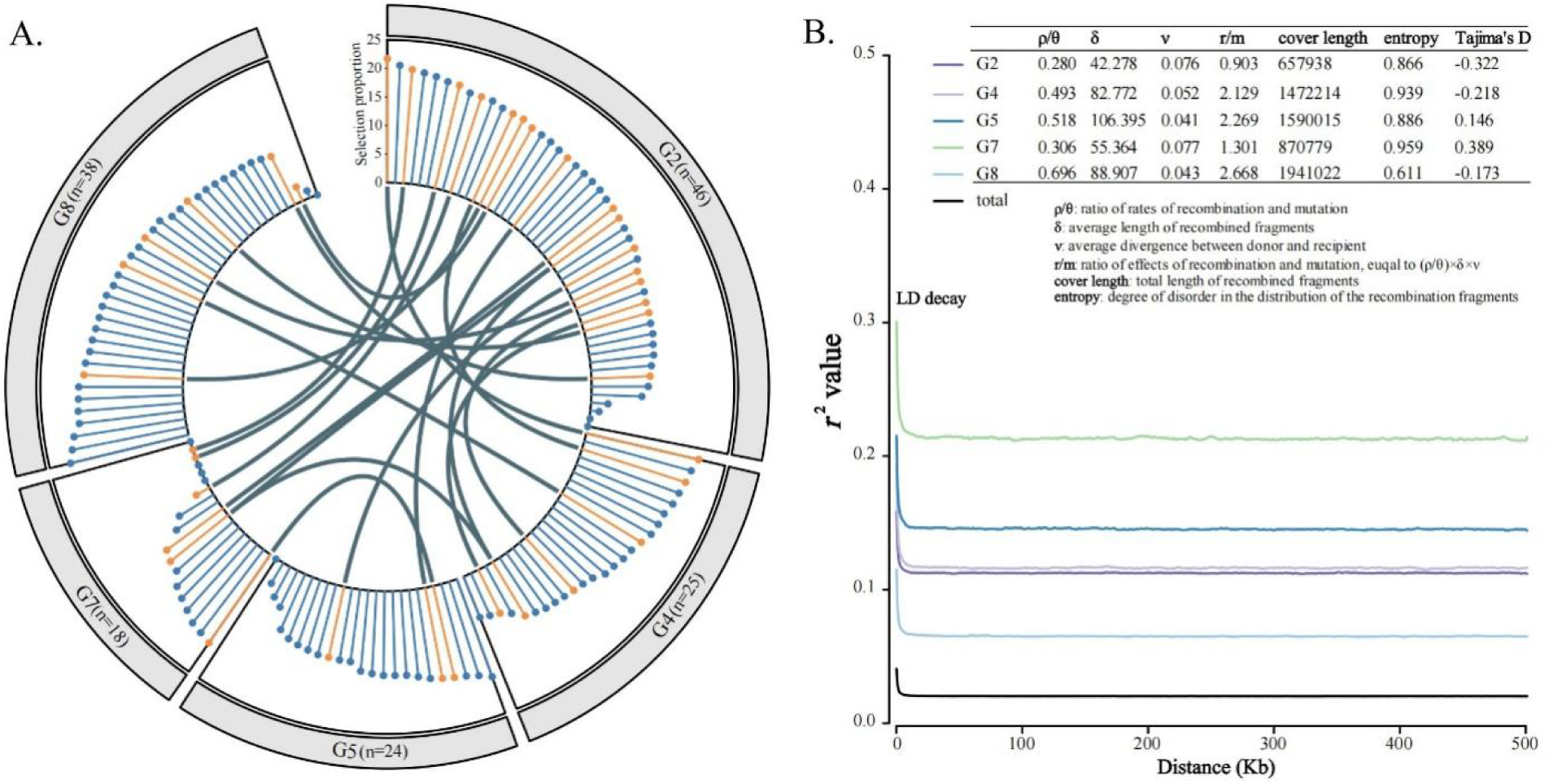
Core genomic evolutionary dynamics and recombination patterns. (**A**) Distribution of genes experiencing positive selection across phylogroups. The *y*-axis represents the proportion of sites exhibiting a Ka/Ks ratio > 1 within individual genes. Genes subjected to positive selection across multiple phylogroups are highlighted in yellow, while inner cyan connecting lines denote instances of potential convergent selection pressures. (**B**) Quantitative parameters characterizing homologous recombination dynamics and population genetic architecture within phylogroups. Key metrics include: ρ/θ, the ratio of recombination to mutation rates; δ, average length of recombined genomic fragments; ν, average sequence divergence between donor and recipient lineages; r/m, the ratio of recombination to mutation effects, calculated as ρ/θ × δ × ν; coverage length, representing the cumulative length of recombined fragments; and entropy, quantifying the distributional disorder of recombination events across the genome. The linkage disequilibrium decay pattern demonstrates the progressive weakening of correlations between genetic loci as chromosomal distance increases, reflecting the influence of recombination on population genetic structure.

Functional characterization of PSGs revealed that G2 possessed a broader functional repertoire (**Table S2**), potentially indicating multi-dimensional selection pressures encountered during evolutionary processes. Although all phylogroups experienced selection in identical metabolism pathways, including cell wall synthesis, damage repair, energy production, signaling, and transport, the specific genes involved differed markedly. These results reflect distinct adaptations to arid microhabitats (Murik, et al. 2017; Li, et al. 2022), enhanced environmental interaction capabilities (Simm, et al. 2015), and the ability to switch to appropriate redox potential strategies in response to extreme environmental fluctuations (Adler, et al. 1999). The “function conserved-gene divergent” pattern likely depicts differentiated optimization strategies employed by various phylogroups while maintaining similar physiological functions, which is congruent with the concept that diverse abiotic pressures can drive adaptive evolution at distinct nodes within the same metabolic pathway (Gonçalves, et al. 2024). Cell wall decomposition and menadione-specific isochorismate synthase (*menF*) pathways underwent co-selection in G2 and G8. However, G2 prioritized N-acetylmuramoyl-L-alanine amidase (*amiABC*), primarily promoting cellular autolysis and division under nutrient-poor conditions (Babu, et al. 2011), whereas G8 favored glycoside hydrolase family 24 (GH24), a lysozyme variant capable of cleaving bacterial cell walls (Kombrink, et al. 2019). Considering the crucial role of *menF* in vitamin K2 synthesis (Daruwala, et al. 1996), it suggests that G2 and G8 acquired survival advantages under oligotrophic and eutrophic conditions, respectively. Additionally, the positive selection of dark-responsive protochlorophyll oxidoreductase (*chlN*) in phylogroup G5 indicates its unique adaptation mechanisms to low-oxygen and dim-light environments (Yamamoto, et al. 2009; Nomata, et al. 2014). In contrast, the selection of circadian rhythm (*kaiC*) and rRNA processing components in G7 suggests a reliance on circadian regulation for resource allocation (Reimers, et al. 2017) or adaptation to hyperosmotic environments (Terrettaz, et al. 2023). The convergent selection observed between G2 and G4/G8 in growth and protein synthesis, between G2 and G5 in damage repair, between G2 and G7 in material transport and energy production, and between G5 and G7 in signal transduction could reflect the prevalent stresses such as resource competition and DNA damage in arid environments. Collectively, the multi-dimensional convergence selection in the core single-copy genes of *M. vaginatus* resembles convergent evolution patterns observed in closely related microorganisms inhabiting extreme environments (Liao, et al. 2024), aligning with the cross-population conservative adaptation paradigms.

Core genome homologous recombination analysis revealed that the recombination-to-mutation rate ratio (ρ/θ) in *M. vaginatus* (**Fig. 2B**) exceeded the threshold characteristic of sexual-like bacterial populations (approx. 0.25) (Fraser, et al. 2007), despite a minimum ANI of 90.3%. It distinguishes *M. vaginatus* from high-ANI clonal *Roseobacter* and sexual-like *Streptomyces olivaceus* (Wang, et al. 2020; Wang, et al. 2022). Contrary to PSG patterns, G2 exhibited the lowest ρ/θ value among phylogroups and a more clonal nature (Shapiro and Polz 2015), while the other lineages followed similar trends to positive selection, with higher values in G8, medium values in G4 and G5, and lower values in G7. Recombination fragment length (δ) was most extended in phylogroup G5, followed by G4 and G8, but shortest in G2 and G7, exhibiting an inverse relationship with the donor-recipient divergence (ν). It supports the recombination termination model, wherein elevated sequence divergence increases the probability of RecA filament dissociation, thus shortening the recombination fragments (Förster, et al. 2023). After accounting for the opposing effects of δ and ν, both the recombination-to-mutation effect ratio (r/m) and genomic coverage by recombination events followed identical trends to ρ/θ. Correspondingly, the proportion of PSGs affected by homologous recombination was higher than 0.9 in G5, G8, and G4 but lower in G2 and G7 (**Table S2**).

Homologous recombination introduces novel mutations while disrupting linkage disequilibrium and selective sweeps (Maynard-Smith and Haigh 1974; Samuk, et al. 2017). As a result, the nucleotide diversity of PSGs (π_pos_) was lower than that of the entire core genome (π_core_) in G2 and G7 but higher in G4 and G5 (**Fig. S11**). In G8, π_pos_ closely approximated π_core_ under high recombination effects (r/m = 2.668), reflecting the dual role of gene flow in restoring local diversity and promoting population homogenization during adaptive evolution (Samuk, et al. 2017; Gonzalez-Torres, et al. 2019). Despite extensive recombination coverage, the entropy analysis revealed that G8 had concentrated recombination hotspots, whereas G7 exhibited shorter and more dispersed patterns of recombination. Focusing on genes experiencing recombination across more than one-third of the evolutionary nodes within populations (**Table S3**), G7 ranked highest (n = 65), followed by G8 (n = 13), G2 (n = 5), G4 (n = 2), and G5 (n = 2). It suggests that the G7 may enhance population adaptability under higher environmental pressure by continuously recombining specific genomic regions to generate adaptive evolutionary sources (Raeside, et al. 2014). It is noteworthy that five genes underwent frequent recombination across all phylogroups, functioning in membrane anchoring, pilus assembly, and protein/DNA regulation, speculating that the species could optimize movement-related surface structures and metabolic regulation to enhance environmental interaction abilities (Averhoff, et al. 2021; Menon, et al. 2021). Therefore, *M. vaginatus* exhibits a sexual-like population structure with distinct recombination patterns among phylogroups, while key recombination genes consistently provide adaptation advantages.

Linkage disequilibrium decay patterns are jointly influenced by natural selection and homologous recombination (Slatkin 2008). The measure *r*^2^ between arbitrary pairwise loci was significantly higher in G7 than in other phylogroups due to its minimal selection and recombination levels. G2, characterized by high selection and low recombination, ranked second, followed by G4 and G5 with moderate selection and recombination. G8, exhibiting high selection and recombination, demonstrated the lowest *r*² values (**Fig. 2B**). Linkage disequilibrium patterns reflect historical population demographic dynamics (Park 2012), wherein a positive value typically indicates population bottlenecks, while a negative value suggests population expansion. Tajima’s D values were negative in G2, G4, and G8 but positive in G5 and G7, further corroborating demographic history inferences and aligning with recent findings on the *Microcoleus* genus (Stanojkovic, et al. 2024). These results indicate that the former three phylogroups experienced expansion while the latter two underwent contraction. G2 primarily adapted to arid habitats through selection and periodic sweeps, whereas G8 adapted to semi-humid habitats through selection and recombination (Shapiro and Polz 2015).

Regarding accessory genomic compartments, the evolutionary history of gene families revealed lineage-specific turnover patterns (**Fig. 3**). The G7 exhibited substantially greater gene turnover than other lineages, with gene loss predominating (gain: 228, loss: 312), which facilitated lower metabolic costs and stronger niche competition (Giovannoni, et al. 2005; Niehus, et al. 2015). Beyond categories with unknown functions and low-proportion complex categories (Other), gene turnover mainly involved categories T (signal transduction), C (energy production), K (transcription), and L (recombination and repair). G4 demonstrated only gene acquisition and expansion, with nearly 60% representing unknown functions, while the remainder primarily belonged to categories T, L, and O (post-translational modification). This pattern may relate to adaptation to complex local environments (Chen, et al. 2021). In G2, gene gains mainly occurred in categories T, N (cell motility), C, and G (carbohydrate metabolism), while gene turnovers involved category O. G8 acquired genes related to T and L, lost a gene involved in category O, and underwent turnover in categories K and H (coenzyme metabolism), suggesting that G2 and G8 adopt life history strategies favoring high growth potential and efficient resource acquisition (Li, et al. 2022). In contrast, G5 exhibited minimal gene family changes, involving only losses in category K and turnover in category O, reflecting its adaptation to low-stress, resource-stable niches.

**Fig. 3.**
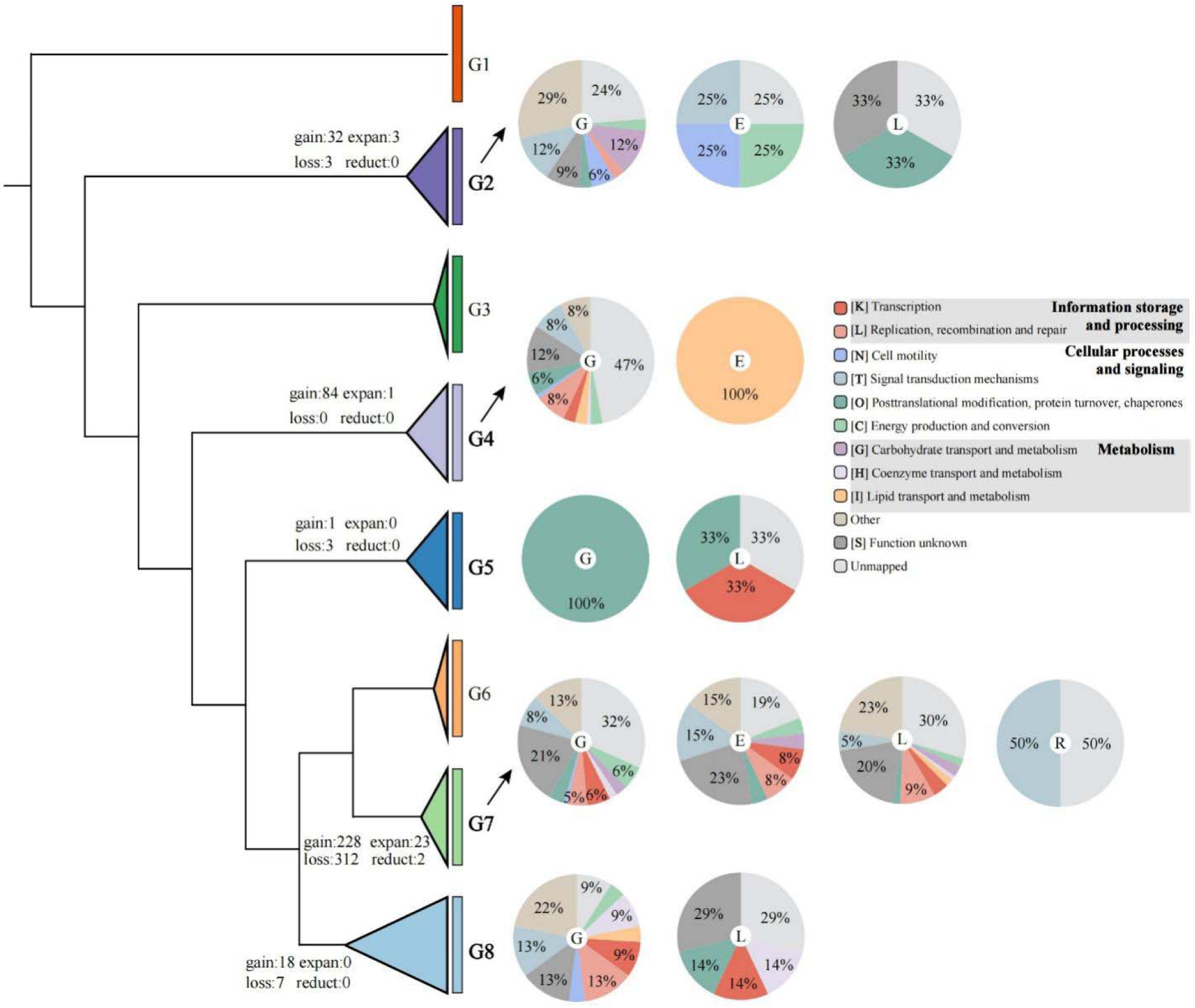
Gene family evolutionary dynamics and functional categorization at phylogroup ancestral nodes. Excluding G1, G3, and G6 due to insufficient sample sizes, the quantitative representation of gene family gain, expansion, loss, and reduction events is annotated adjacent to each ancestral node, illustrating the dynamic genomic content transformations during lineage differentiation. Gain events are defined as instances where gene families transition from absence in parental nodes to single-copy presence in target nodes; conversely, loss events represent the elimination of gene families that existed as single-copy in parental nodes. Expansion events denote transitions from single-copy to multiple-copy states between parental and target nodes, while reduction events indicate the reverse trajectory from multiple-copy to single-copy configurations. Accompanying pie charts illustrate the proportional distribution of functional categories for each evolutionary event type, with categories representing <10% across all nodes consolidated under “Other”. Central letters within pie charts designate specific event types: **G**, gene family gains; **E**, expansions; **L**, losses; **R**, reductions. The analysis reveals the functional basis underlying adaptive genomic restructuring processes across distinct phylogroups.

Genome size is influenced by gene gain and loss dynamics and positively correlates with effective population size in bacteria (Bobay and Ochman 2017b). The ability of bacterial species to capture and maintain a diverse accessory gene pool likely enables them to occupy varied environments and sustain large population sizes. Contrary to previous reports suggesting that most bacterial genomes tend to shrink due to deletion mutations driven by genetic drift (Bobay and Ochman 2017b), our results showed a gradual increase in gene family numbers along the phylogeny (**Fig. S12**). The expansion rate in G4 is the lowest, consistent with its reduced HGT frequency (**Fig. S9B**) and constraints on genome expansion imposed by nutritional limitations (Bobay and Ochman 2017b), drought stress (Schley, et al. 2022), and salinity (Dong, et al. 2024).

The observed pattern of genome size change contradicts the expected positive correlation with effective population size (Bobay and Ochman 2017b), highlighting the prominent contribution of natural selection over genetic drift in evolution. These findings suggest that *M. vaginatus* genomes have undergone gradual expansion under natural selection to adapt to variable environments, while developing diverse life history strategies that range from high competition to stable colonization through the evolution of accessory genomes. Signal transduction and its downstream responsive genes constitute the core of adaptation strategies across the eight phylogroups.

### GWAS identifies differentiation-associated loci

To comprehensively screen differentiation-associated variants across the entire genome, we conducted a genome-wide association study (GWAS) targeting population divergence patterns. Our analysis identified 737 single-nucleotide polymorphisms (SNPs) that were significantly correlated with divergence (*p* < 5×10^-8^; **Fig. 4A**), with statistical model validity confirmed by quantile-quantile (Q-Q) plot analysis (**Fig. S13**). These loci were distributed throughout the genomic landscape, mapping to 270 core genes (100% prevalence), 55 soft-core genes (95% to 99%), 42 shell genes (15% to 95%), and eight cloud genes (1% to 15%), consistent with the core genome’s predominant contribution to cryptic speciation processes (Wang, et al. 2020). The SNPs formed multiple linkage disequilibrium blocks across the genome (**Fig. 4B**). Among these differentiation-associated variants, only 28 SNPs corresponded to previously identified evolutionary signatures, with 16 overlapping with positive selection signals and the other 12 associated with gene gain or loss events (**Table S4**). Among the 709 unmatched loci, approximately 54% (381 sites) demonstrated linkage (*r*^²^> 0.6) with matched sites, while nearly half represented neutral sites that lacked hitch-hiking effects (Maynard-Smith and Haigh 1974).

**Fig. 4.**
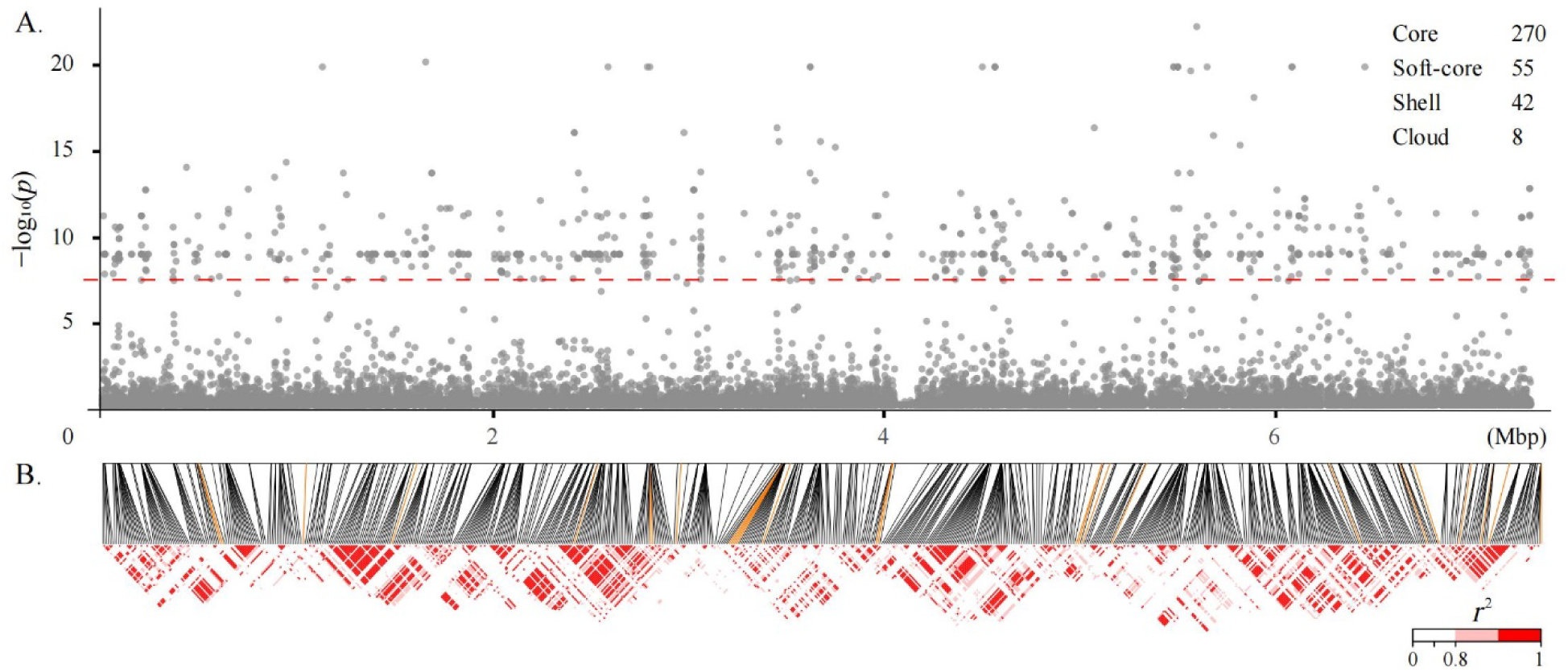
Genome-wide association analysis reveals differentiation-associated genetic variants. (**A**) Manhattan plot displaying the statistical significance of genome-wide SNPs across chromosomal positions. The red dashed line represents the significance threshold (*p* < 5 × 10^-^), with data points exceeding this threshold considered significantly associated with phylogroup differentiation. The accompanying table categorizes genes harboring significant SNPs according to genomic prevalence: core genes (present in 100% of genomes), soft-core genes (present in 95% to 99% of genomes), shell genes (present in 15% to 95% of genomes), and cloud genes (present in multiple genomes but < 15% of total genomes). (**B**) Linkage disequilibrium architecture among significant loci, where color intensity gradient from white to red represents escalating LD strength, with only *r*^2^ > 0.8 indicating functionally linked loci. Orange directional lines identify loci corresponding to genes previously implicated in evolutionary processes, specifically positive selection and gene family turnover events, thereby connecting population-level genetic associations with established evolutionary mechanisms.

This pattern supports the theoretical framework in which adaptive fitness enhancement involves a shift from positive selection to neutral evolutionary processes (Kishimoto, et al. 2010). Notably, phylogroups G2 and G7 exhibited more substantial hitch-hiking effects around evolutionary event-associated SNPs, corresponding to the abovementioned low recombination rates (**Fig. 2B**). Functional characterization revealed that most genes harboring the 28 matched SNPs are still unknown, while the remainder participated in various cellular processes like signal reception and response mechanisms (e.g., chemotaxis), material transport systems (Mg^2+^ and vitamin B12), and protein folding pathways. These findings elucidated that natural selection on the core genome constitutes the predominant driver of *M. vaginatus* population differentiation. This divergence process is initiated by selection and mediated by post-selection neutral evolution and recombination mechanisms (Chen, et al. 2021; Li, et al. 2022), ultimately manifesting as strategic shifts in life history paradigms across the phylogroups.

### Environment shapes intraspecific genetic divergence and evolutionary strategies

To identify environmental drivers underlying genetic subdivision among populations, we collected environmental parameters from each site where *M. vaginatus* strains were sampled (**Table S5**). We then performed distance-based redundancy analysis (dbRDA) to evaluate the explanatory power of key variables on multiple genomic distance metrics: whole genome divergence (WGD), orthologous gene divergence (OGD), core single-copy gene sequence divergence (CGD), and core single-copy protein divergence (CPD) (**Fig. 5A**). The first two ordination axes explained over 65% of the variation for all metrics, indicating robust multivariate ordination performance. We found that phylogroup G7 exhibits significant distinctions from other groups in the measured characteristics, despite the evident divergences among groups being observable. For WGD-based ordination, all phylogroups were more distinctly separated. In contrast, for OGD, CGD, and CPD analyses, a clear separation was observed in G2, G7, and G8, while the other phylogroups exhibited partial overlap. It suggests that environmental impacts on non-coding sequence divergence may resemble the responsive activation of transposable elements (Miller, et al. 2021).

**Fig. 5.**
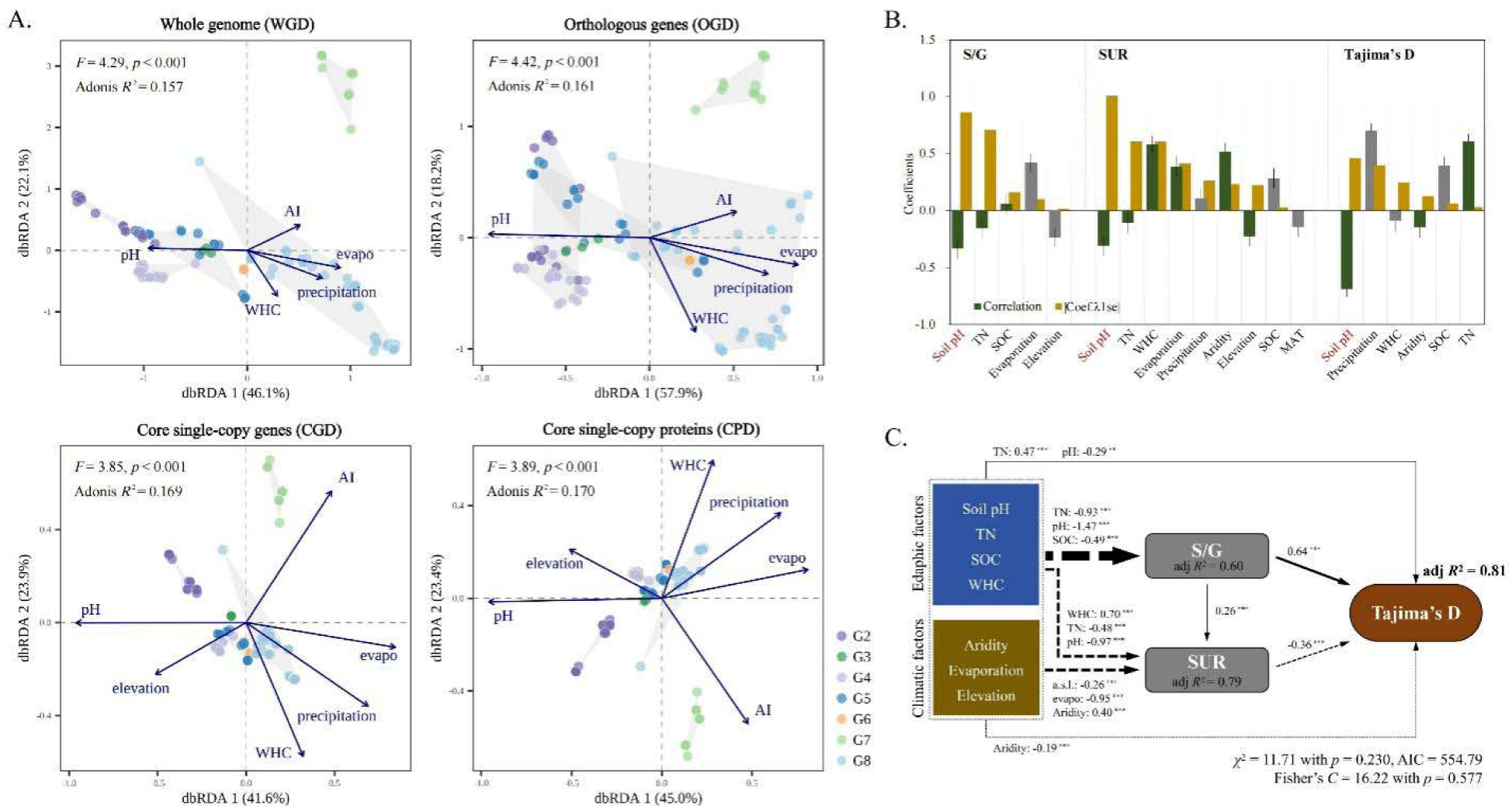
Environmental predictors of intraspecific genetic characteristics and the regulation of evolutionary strategy. (**A**) Distance-based redundancy analysis (dbRDA) examining the relationships between environmental variables and multiple genetic distance metrics: whole-genome distance (WGD), Bray-Curtis distance of orthologous group composition (OGD), maximum likelihood distance of core single-copy genes (CGD), and maximum likelihood distance of core single-copy proteins (CPD). Scatter points are differentiated by phylogroup affiliation, with all labeled environmental variables demonstrating significant effects with moderate collinearity. (**B**) Variable importance assessment and individual correlations with key evolutionary parameters [selection-to-gene gain ratio (S/G), proportion of selected genes undergoing recombination (SUR), and Tajima’s D], evaluated through LASSO regression analysis. Gray histograms represent non-significant factors excluded by *d*-separation tests in subsequent piecewise structural equation modeling (pSEM). (**C**) Causal pathways linking environmental factors to evolutionary strategies, elucidated through the pSEM. Directional arrows represent hypothesized causal relationships, with the orientation of the arrows indicating the direction of influence between variables. Numerical values adjacent to arrows denote standardized path coefficients, where solid arrows represent positive effects and dashed arrows indicate negative effects; arrow width corresponds to effect size. Asterisks denote statistical significance levels (** *p* < 0.01, *** *p* < 0.001). Adjusted *R*^2^ values quantify the proportion of variance explained for each dependent variable. Model adequacy was assessed using Fisher’s *C* statistic and the χ² test, confirming a parsimonious model fit with a corresponding AIC value provided for model comparison.

Key environmental variables exhibited clade-specific effects, including pH, aridity index, precipitation, evaporation, water-holding capacity (WHC), and elevation (**Fig. 5A**). Moisture-related parameters, as precipitation, evaporation, and WHC, predominantly influenced the G8. In contrast, the aridity index had a positive influence on the G7, while the G2 was primarily affected by soil pH. The triangular distribution pattern formed by G2, G7, and G8 in ordination space reflects the integrated effects of these abiotic factors. Soil pH determines the availability of inorganic carbon to cyanobacteria and serves as a modulator of the redox state (Wang and Kuzyakov 2024). In addition, WHC indicates the potential activity status of soil microbes in drylands. The pronounced influence on the G8 phylogroup from semi-humid habitats suggests that strain activity varies with WHC and moisture fluctuations, while genomic convergence of G2 in pH adaptation likely stems from similarities in HCO ^-^utilization strategies (Ramoneda, et al. 2023). For the phylogroup G7, the combined effects of aridity could primarily manifest through frequent cycles of drying and hydration in coastal soils of humid areas. Its significant impact on the G7 indicates that the moisture stress, which is distinct from that in inland arid regions, contributes to codon usage preferences (Hellweger, et al. 2018; Chuckran, et al. 2023).

Precipitation, evaporation, and WHC collectively reflect water availability in soils, while pH governs photosynthetic substrate availability and metabolic activity states. Their combined influence results in the sharp clustering of G2 (Asian arid biocrusts), G7 (humid zones), and G8 (semi-humid multiple habitats), highlighting minimal differences in corresponding metabolic activities within habitats of each phylogroup, but substantial differences between phylogroups. Therefore, local pH levels and the moisture-related variables of climatic zones constitute the most critical macro- and micro-scale factors driving population divergence, similar to the early speciation and rapid evolution of soil bacteria (Chase, et al. 2021; Maya-Lastra, et al. 2024).

To uncover cascading relationships between environmental factors and evolutionary strategies of *M. vaginatus* continuum, we applied piecewise structural equation linear model (pSEM) to analyze the effects of environmental variables on S/G, SUR, and Tajima’s D. We proposed the S/G ratio to represent the proportion of PSGs relative to the number of gene gains, aiming to quantify the relative contributions of different genomic regions to population adaptation. Considering that gene acquisition also reflects natural selection, the S/G ratio can reveal the preferential regions of selective pressure during species divergence. The SUR metric represents the proportion of PSGs that exhibit recombination signals. An increase in SUR indicates that a greater number of selective advantages are disseminated and fixed within populations through recombination, highlighting recombination as a crucial mechanism for the spread of adaptive variants and its important role in facilitating the rapid adaptation and dispersal of beneficial alleles within populations. Given the multicollinearity problem, strongly correlated factors were stepwise eliminated using least absolute shrinkage and selection operator (LASSO) regression (**Fig. 5B**). For abiotic predictors [aridity, evaporation, elevation, soil pH, total nitrogen (TN), and soil organic carbon (SOC)], we found climatic and edaphic variables could directly mediate Tajima’s D but offset with each other (summed net effect size = 0.01, Fisher’s *C* = 16.22 with *p* = 0.577; **Fig. 5C**). In contrast, edaphic indirect influences through S/G ratio (|coefficient| on S/G: 2.92 and S/G → Tajima’s D: 0.64, *p*-values < 0.001) dominate the cascading pathways, rather than climatic effects (|coefficient| on SUR: 0.81 and SUR → Tajima’s D: -0.36). The environmental predictors exhibited consistent negative effects on S/G and SUR, with contrasting subsequent effects on Tajima’s D, highlighting the trade-off between evolutionary strategies in *M. vaginatus* strains. The results demonstrated that local pH condition serves as the primary driver of population divergence, while climatic variables (aridity, evaporation, and elevation) moderately mediate shifts in historical evolutionary signals (**Fig. 6**). In these strategic adjustments, despite the strong effects of microscale edaphic variables (especially pH), SUR and S/G constitute more dominant pathways that directly promote changes in Tajima’s D, serving as the core mechanism that reflects the balance between selection pressures and genetic stability during the divergent evolutionary processes of this cosmopolitan cyanobacterium.

**Fig. 6.**
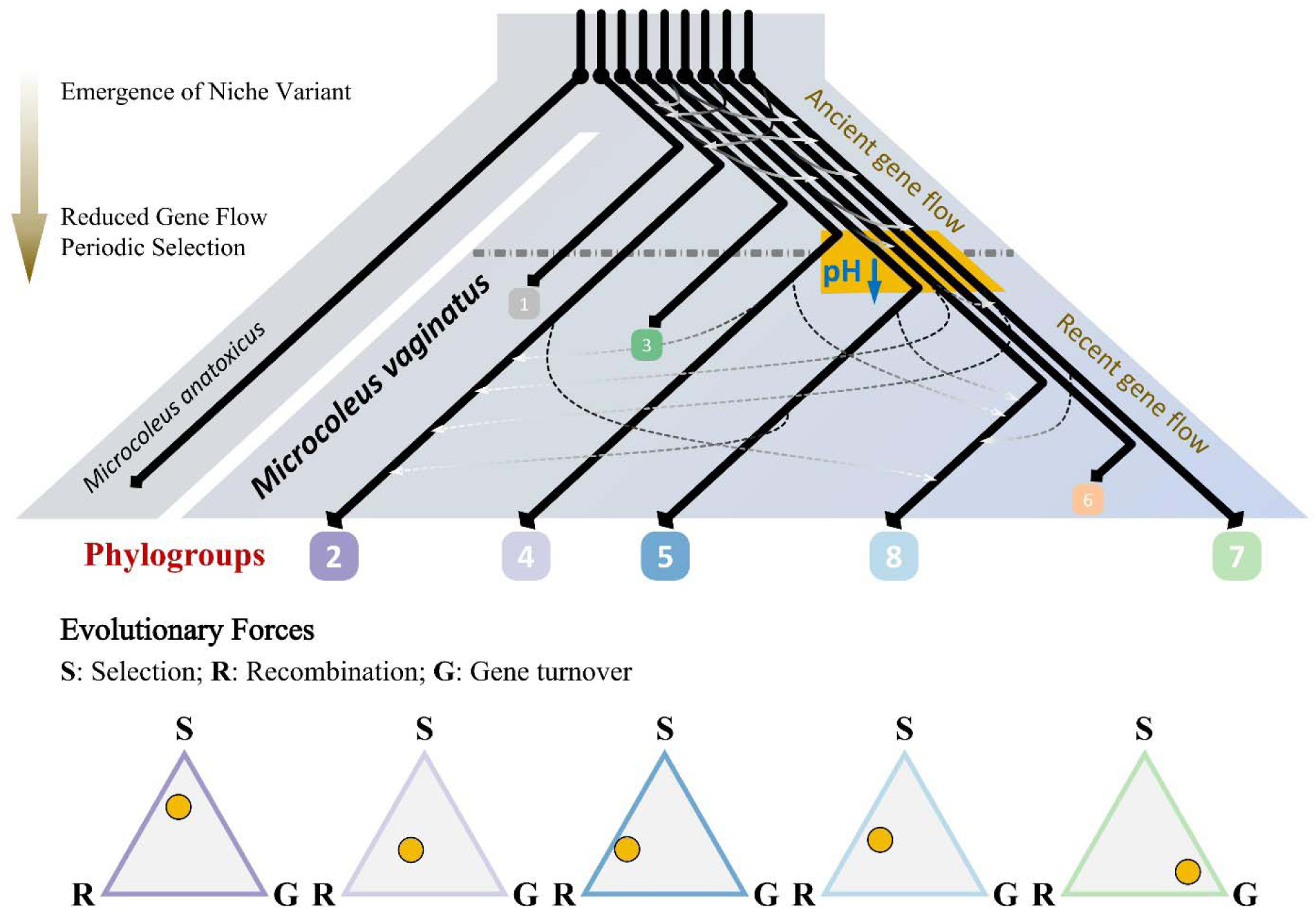
Conceptual model of divergence dynamics in the cyanobacterial species *Microcoleus vaginatus*. The upper panel depicts the topological architecture of differentiation processes, organized according to the maximum likelihood phylogenetic tree constructed from concatenated single-copy core protein sequences. With the establishment of gene flow barriers with the outgroup *Microcoleus anatoxicus*, phylogroups experienced extensive ancient gene flow despite the emergence of ecological niche variants. Subsequent reduction in recent gene flow, coupled with periodic selection events, facilitated phylogroup separation. The horizontal gray line indicates the temporal boundary between ancient and recent evolutionary phases, with five dominant phylogroups distinguished. Through reconstruction of ancestral node environmental conditions, pH played a critical role in the tipping point separating G4 from the subsequent phylogroups (G5-G8), marking a transition from alkaline to acidic conditions across this divergence event (data not shown). The lower panel illustrates the relative contributions of key evolutionary forces during phylogroup divergence, with each triangular plot quantifying the proportional importance of these mechanisms across different phylogroups. Abbreviations: **S**, positive selection; **R**, homologous recombination; **G**, gene family turnover.

To conclude, based on a comprehensive whole-genome analysis of 132 global *Microcoleus vaginatus* strains, this study reveals eight distinct monophyletic lineages exhibiting remarkable intraspecific genetic plasticity with substantial gene flow, providing compelling evidence for ongoing speciation processes. Population divergence emerges through dynamic equilibrium among positive selection, homologous recombination, and gene family turnover, with complementary evolutionary patterns manifesting between core and accessory genomic compartments (**Fig. 6**). Our findings demonstrate that adaptive evolution primarily involves strategic shifts in life-history paradigms driven by environmental pressures, wherein diverse abiotic stresses can drive evolution at distinct nodes within identical metabolic pathways.

Environmental factors, particularly pH and aridity, play pivotal roles in shaping genetic architecture and evolutionary trajectories. A set of abiotic variables mediates complex regulatory mechanisms that govern evolutionary strategies. These multifaceted processes collectively promote the diversification of adaptive strategies and facilitate efficient niche colonization across heterogeneous environments. Ultimately, *M. vaginatus* thrives and exhibits rapid evolutionary responses across diverse ecological contexts through sophisticated trade-offs among multiple evolutionary mechanisms. This study sheds light on the intricate evolutionary dynamics of non-model microorganisms and elucidates the environment-driven mechanisms that underpin evolutionary adaptations in globally distributed prokaryotes.

## Materials and Methods

### Sample collection, cyanobacterial cultivation, and genome sequencing

We collected biocrust samples from arid and semi-arid regions spanning a 3,400 km transect across northern China. Benthic cyanobacterial mats were also sampled from diverse aquatic environments, including open channels, wetlands, and temporary puddles (**Table S1**). Samples were cultivated on BG11 medium plates under continuous illumination, and emerging cyanobacterial colonies were subsequently isolated and purified through a standard pipeline. A total of 52 *M. vaginatus* strains were identified through microscopy examination and full-length *rrs* alignment. Besides, five additional strains were obtained from FACHB to supplement our collection.

Genomic DNA was extracted from each strain using the DNeasy Plant Mini Kit (Qiagen GmbH, Germany), with modifications based on the CTAB extraction protocol. Six samples (strains APE-1 to APE-6) were subjected to paired-end (PE) 150-bp sequencing on the Illumina HiSeq Xten platform. The other samples underwent dual-platform sequencing, employing second-generation Illumina PE 150-bp sequencing kits on the NovaSeq 6000 platform and third-generation Oxford Nanopore PromethION technology, which generated over 11 GB of raw sequencing data. Following quality filtration to remove low-quality reads, long-read sequences were assembled using Flye v2.9.1 (Kolmogorov, et al. 2019). Based on short-read data integration, the resulting assemblies were subsequently polished with Pilon v1.23 (Walker, et al. 2014). Genomic contigs were binned using metaWRAP v1.3.2 (Uritskiy, et al. 2018), and the quality of each assembled genome bin was evaluated using CheckM to assess assembly completeness and contamination (Parks, et al. 2015).

### Global dataset construction, genome annotation, and phylogenetic reconstruction

A comprehensive global dataset was assembled by incorporating 75 high-quality genomes (completeness > 95%, contamination < 5%) harboring the diagnostic 11-bp conserved insertion (5’-GCA ACC TGA CG-3’) within the *rrs* fragment (Garcia-Pichel, et al. 2001), which were retrieved from public databases and integrated into our analytical framework (**Table S1**). These genomes encompassed strains originating from North America, Africa, and Europe, thereby providing broad biogeographic representation. Environmental factors (mean annual precipitation, temperature, evaporation, aridity index, windspeed, elevation, TN, pH value, and SOC) corresponding to individual strains were extracted from established public databases, including WorldClim (http://worldclim.org/), SoilGrids (https://soilgrids.org/), and the Global Aridity Index and Potential Evapotranspiration (ET0) Database (Trabucco and Zomer 2019), utilizing the geographic coordinates of their respective isolation sites. The WHC parameter for each terrestrial strain was estimated based on the clay content of soils, while the WHC for aquatic strains was intuitively assigned as 100% (Clapp and Hornberger 1978).

Ribosomal RNA gene sequences were predicted for each *M. vaginatus* genome using RNAmmer v1.2 (Lagesen, et al. 2007). Functional genome annotation was performed using DIAMOND v2.0.8.146 and eggNOG-mapper v2.1.8 (https://github.com/eggnogdb/eggnog-mapper) against both the KEGG and eggNOG5 databases (Buchfink, et al. 2015), invoking stringent thresholds (identity ≥ 40%, coverage ≥ 50%) to ensure annotation reliability. Codon usage bias was analyzed using CodonW v1.3, and maximum growth rates were subsequently estimated using the *R* package *gRodon* (Weissman, et al. 2021). Orthologous group clustering was performed using OrthoFinder v2.2.7 (Emms and Kelly 2019), which employs *diamond* for sequence alignment procedures and *msa* for gene tree inference algorithms. Multiple sequence alignments of nucleotide and amino acid sequences of core single-copy genes and *rrs* were conducted using MAFFT v7.453 with optimized parameters (*-localpair* and *-maxiterate* = 1000) (Katoh and Standley 2013). Alignments were then trimmed using TrimAl v1.4.rev22 with the parameter *-automated1* to remove poorly aligned regions (Capella-Gutierrez, et al. 2009).

Phylogenetic trees based on concatenated trimmed alignments were reconstructed using IQ-TREE v1.6.11 with the ultrafast bootstrap method implemented within the IQ-TREE framework (Nguyen, et al. 2015; Hoang, et al. 2017). Optimal substitution models were determined using the built-in ModelFinder algorithm (Kalyaanamoorthy, et al. 2017), yielding the following best-fit models: JTT+F+I+G4 for core single-copy gene proteins, and SYM+I+G4 for core single-copy gene nucleotides and rRNA sequences. Population structure was further validated using an improved hierarchical Bayesian clustering algorithm implemented in the *R* package *fastBAPS*. Additionally, we reconstructed a phylogenetic network using SplitsTree4 v4.17.1 to visualize reticulate evolutionary relationships (Huson and Bryant 2006).

### Intraspecific diversity and evolutionary event estimation

Genetic diversity assessment was conducted through pairwise BLAST alignment of *rrs*, complemented by calculation of pairwise ANI across all 132 genomes using fastANI v1.2 with optimized fragment length parameter (*-fragLen* = 1000) (Jain, et al. 2018). Given the unavailability of raw sequencing reads for database-sourced genomes, haplotype variant calling was performed against the reference genome from strain APE-401 using Snippy v3.1, which fragmented the provided contigs into 250-bp single-end reads. Following the segregation of complex-type variants, all variant call format files were consolidated into a unified file containing SNP positions across all genomes. Population genetic parameters, including π and Fst, were computed using VCFtools v0.1.16 across 10-kb sliding windows (Danecek, et al. 2011). Tajima’s D statistic was calculated based on core genome alignments to infer demographic history patterns (Tajima 1989).

We calculated h/m ratios across increasing genome samples using ConSpeciFix v1.3.0 to quantify the extent of gene flow. Inter-phylogroup gene flow dynamics were evaluated using the introgression score as previously described (Diop, et al. 2022). DNA sequence similarity assessments were conducted within non-overlapping 100-bp windows spanning the core genome. A genomic fragment was classified as having experienced inter-phylogroup gene flow when at least one genome from the target phylogroup (target) exhibited greater similarity to genomes from alternative phylogroups (source) than the rest of the genomes within its phylogroup.

Differential sequence identity thresholds were applied between source and target phylogroups, with 100% and 95% identity representing recent and ancient gene flow events, respectively. HGT events between phylogroups were also detected using MetaCHIP v1.10.13 (Song, et al. 2019).

Evolutionary selection pressures were quantified by calculating Ka/Ks for core genes using the *codeml* module in PAML v4.9 under branch-site models(Yang 2007). Likelihood ratio χ² tests were used to compare the null and alternative models, with a significance threshold of 0.01 applied to identify genes containing sites with a Ka/Ks ratio greater than 1, indicating positive selection. Homologous recombination was estimated using ClonalFrameML v1.12 (Didelot and Wilson 2015), and the entropy of recombinant fragments was calculated following the Shannon entropy formula:

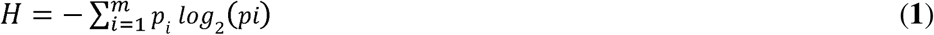

where *p_i_* denotes the frequency of recombination fragments distributed across the core genome. Linkage disequilibrium decay patterns were analyzed using PopLDdecay v3.43 with a maximum SNP distance threshold of 500 kb (Zhang, et al. 2019). Visualization was achieved through the Perl script Plot_MultiPop.pl provided within the software package, by employing the parameters*-bin1* = 100 and *-bin2* = 1000 for optimal resolution. Gene gain and loss events at individual phylogenetic nodes, along with their associated posterior probabilities, were inferred using COUNT (Csuros and Miklos 2009). The phylogenetic birth-death model utilized a discrete γ distribution with categories configured as length:gain:loss:duplication = 4:1:1:4, with the maximum number of optimization iterations set to 100 to ensure convergence.

### Genome-wide association assessment

We implemented a quality control pipeline for our genome-wide association analysis. Firstly, we excluded SNPs with minor allele frequency ≤ 0.05 and those located within five base pairs of insertion-deletion polymorphisms. The filtered dataset was then used to construct a genetic relationship matrix, and associations were estimated using a mixed linear model implemented in GEMMA. To ensure robust statistical inference, we employed three complementary tests: Wald, likelihood ratio, and score tests. We selected the most significant *p*-value for each locus among these tests and applied the Benjamini-Hochberg method for multiple testing corrections. We established a significance threshold of 5 × 10^-8^ for identifying significant associations. The results were visualized using the Manhattan function from the *R* package *qqman*, while the consistency of our findings was assessed through a Q-Q plot comparing observed and expected *p*-values. In addition, we characterized linkage disequilibrium blocks among significant loci using the *R* package *LDheatmap*.

### Statistical analyses

All statistical analyses were performed using *R* version 4.0.2, with other software specified as needed. We calculated four genetic pairwise distance metrics for dbRDA: WGD as (1 - ANI)/10, OGD using Bray-Curtis distances computed with the vegdist function from the vegan package, and both CGD and CPD from maximum likelihood distances obtained during phylogenetic reconstruction with IQ-TREE. A robust random forest imputation method was employed to predict missing values in the environmental matrix using the *missForest* package. The dbRDA analysis was performed using the *dbrda* function from the *vegan* package. To assess the independent explanatory power and statistical significance of environmental factors, we employed the *rdacca.hp* and *envfit* functions, respectively. We evaluated the pairwise correlations between environmental variables using Spearman’s rank correlation method. Ultimately, we constructed pSEMs using the *piecewiseSEM* package to elucidate the cascading relationships among environmental variables, the S/G ratio, SUR 6, and the population history signal (i.e., Tajima’s D).

## Acknowledgments

We thank Zuowen Wang and Xiaoyu Guo for their contribution to the isolation of *M. vaginatus* strains and FACHB for providing strain collections. We also appreciate Chengcai Zhang for his constructive suggestions on the manuscript draft. This study is supported by the National Natural Science Foundation of China (grant No. 32370125 and 32430005), Natural Science Foundation for Distinguished Young Scholars of Hubei Province (grant No. 2022CFA105), and Strategic Priority Research Program of the Chinese Academy of Sciences (grant No. XDA17010502).

## Data and code availability

The genomes reported in this study are publicly available from the NCBI BioProject database under the accession number PRJNA1052756. The scripts supporting the findings in this study are deposited on GitHub (https://github.com/rosemed/Microcoleus-vaginatus-evolution). The raw climatic data are obtained from the WorldClim (http://worldclim.org/), SoilGrids (https://soilgrids.org/), and the Global Aridity Index and Potential Evapotranspiration (ET0) Database (https://doi.org/10.6084/m9.figshare.7504448.v2).

## Author contributions

Conceptualization: HL, CH; Methodology: JW, HL, CH; Investigation: JW, HY; Visualization: JW, HL; Supervision: RL, WS, CH; Writing—original draft: JW, HL; Writing—review & editing: HL, RL, WS, CH.

## Supplementary Materials

The manuscript is accompanied by Supplementary Materials with 13 Figures and five Tables.

