## Supplementary Materials for "pH-driven evolutionary divergence and ongoing speciation in the cosmopolitan cyanobacterium *Microcoleus vaginatus*"

**Supplemental materials** contain 13 Figures and five extended Tables.

**Figure S1.** Comparison between 16S rRNA gene (*rrs*) similarity and average nucleotide identity (ANI).

**Figure S2.** The relationship between minimal doubling time (d, in days) and 16S rRNA gene (*rrs*) copy number across strains.

**Figure S3.** Phylogenetic architecture with outgroup.

**Figure S4.** Comparison between the phylogenetic tree and fastBAPS clustering.

**Figure S5.** Phylogenetic relationships and genomic codon usage variation among phylogroups.

**Figure S6.** Phylogenetic network among phylogroups.

**Figure S7.** Phylogenetic tree based on 16S rRNA gene (*rrs*) sequences.

**Figure S8.** Inconsistency of species delimitation based on different thresholds.

**Figure S9.** Genomic exchange among phylogroups.

**Figure S10**. Phylogenetic architecture excluding positively selected genes (PSGs).

**Figure S11.** Normalized density distribution of nucleotide diversity within genomic regions under positive selection and the total core genome.

**Figure** **S12**. Relationship between gene family count and phylogenetic distance from the ancestral node across evolutionary nodes within each phylogroup.

**Figure S13.** Quantile-Quantile (Q-Q) plot of genome-wide association study (GWAS).

**Figure S1.** Comparison between 16S rRNA gene (*rrs*) similarity and average nucleotide identity (ANI). Normalized density distribution of pairwise *rrs* similarities (pink) and ANI values (blue). All *rrs* sequences were extracted from whole-genome sequences using RNAmmer v1.2 and subjected to BLASTn alignment to obtain pairwise similarities. Pairwise ANI values were calculated by fastANI v1.2 with optimized fragment length set as 1000, where overlap rates of fragments between any pair of genomes exceeded 60%. Some non-circularized, contig-level genomes harbored multiple *rrs* sequences with substantial sequence divergence, resulting in a secondary peak at approximately 80% in the *rrs* similarity distribution.

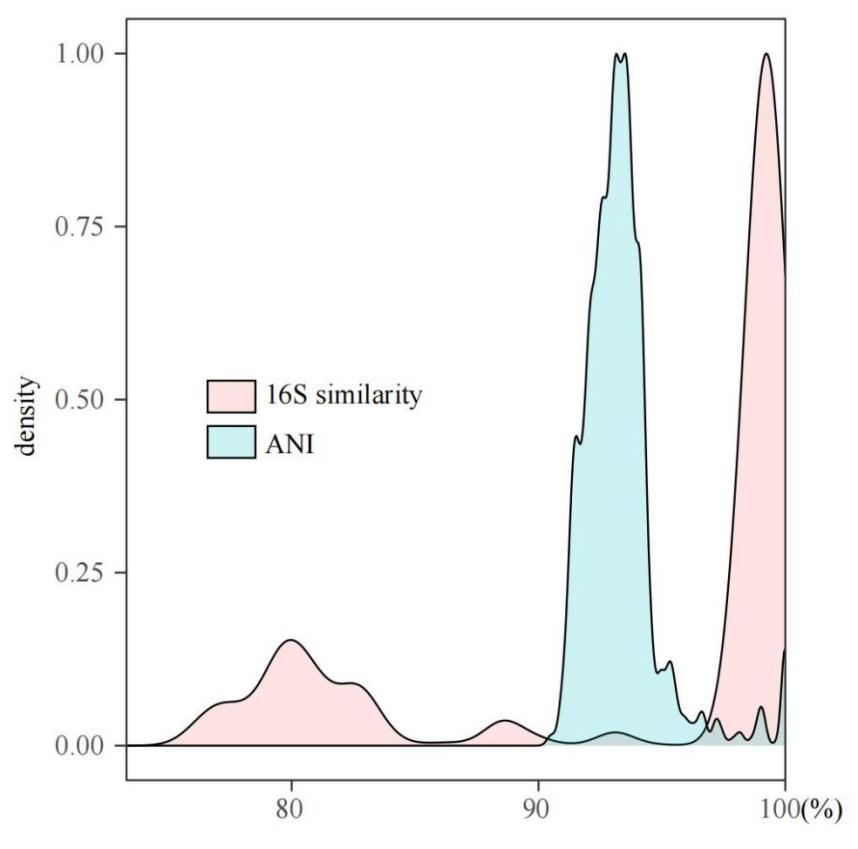

**Figure S2.** The relationship between minimal doubling time (d, in days) and 16S rRNA gene (*rrs*) copy number across strains. The d values corresponding to each genome were estimated using the *R* package *gRodon* based on codon usage bias within the genomes. To avoid underestimating the number of *rrs* sequences within a genome due to sequence fragmentation, only circularized genomes were considered. Contrary to the previous understanding, a higher number of *rrs* sequences does not correspond to a lower d value, with no significant correlation observed between the two variables.

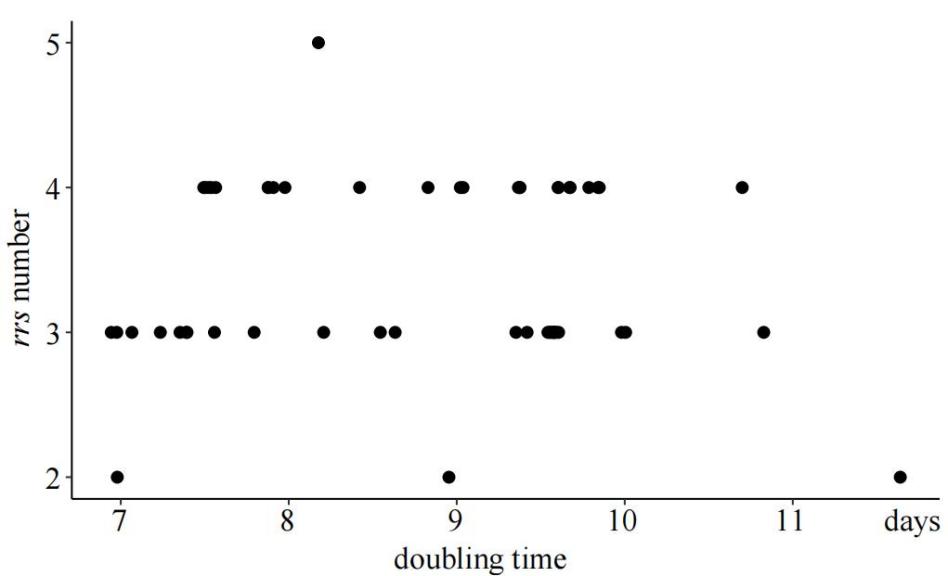

**Figure S3.** Phylogenetic architecture with outgroup. A circular phylogenetic tree was constructed using concatenated core single-copy protein sequences to infer evolutionary relationships among the strains. *Microcoleus anatoxicus* CHAB4116 was used as the designated outgroup to root the tree. Branch lengths are proportional to the number of substitutions per site, as indicated by the scale bar (0.05 substitutions/site). Node support was evaluated using bootstrap analysis, with highly supported nodes (bootstrap values ≥ 75) marked in red to highlight robustness of the inferred relationships. The topology reveals that the tree containing only *Microcoleus vaginatus* strains should be re-rooted at the parent node of strain FACHB1854.

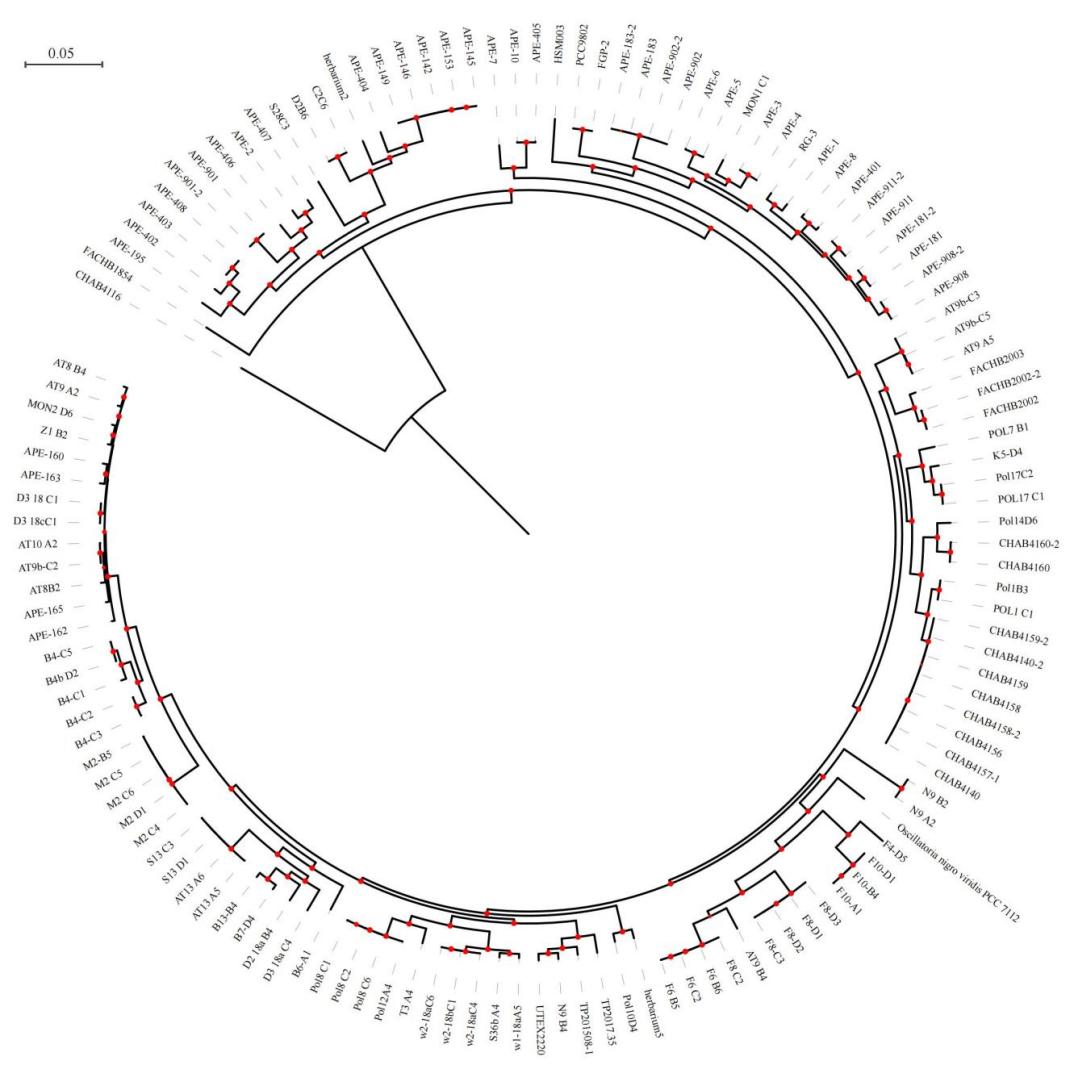

**Figure S4.** Comparison between the phylogenetic tree and fastBAPS clustering. The left panel displays the maximum likelihood phylogenetic tree constructed from concatenated core single-copy protein sequences. The right panel presents a heatmap generated from fastBAPS clustering results. This Bayesian hierarchical clustering method groups genomes based on an approximate fit to a Dirichlet process mixture model for clustering multi-locus genotype data. Darker shades of blue blocks indicate tighter clustering between samples. The arrangement of samples in the heatmap corresponds to the ordering of taxa on the phylogenetic tree, allowing direct comparison between phylogenetic inference and clustering patterns. The middle color blocks correspond to monophyletic phylogroups G1 to G8.

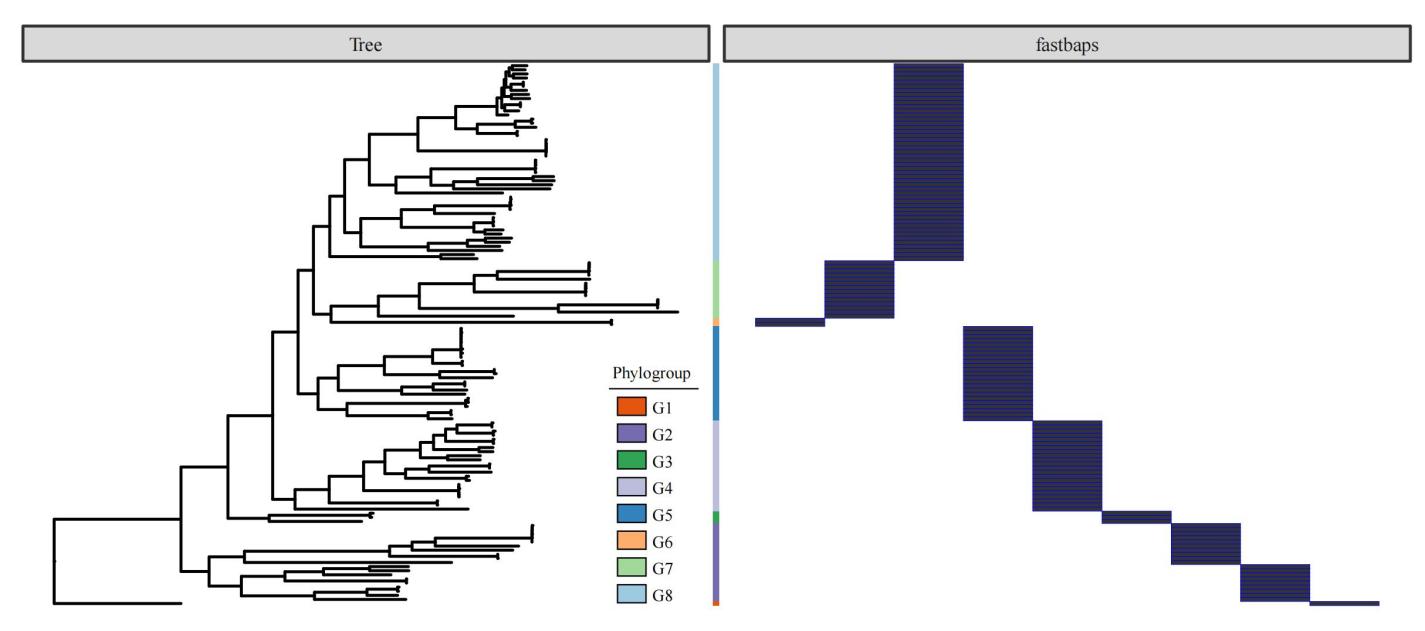

**Figure S5.** Phylogenetic relationships and genomic codon usage variation among phylogroups. (A) A maximum likelihood phylogenetic tree was reconstructed from concatenated nucleotide sequences of core single-copy genes across all sampled genomes. Branches are color-coded to indicate membership in one of eight distinct phylogenetic groups (G1 to G8), as shown in the legend. Red dots on the internal nodes represent bootstrap values greater than 75%, indicating strong statistical support for these clades. The scale bar denotes the number of substitutions per site. (B) Principal component analysis of genomic codon usage frequencies calculated from each genome, capturing variation in codon preference patterns among the samples. The two axes represent the first two principal components, PC1 and PC2, which explain 52.67% and 20.71% of the total variance, respectively (values in parentheses). Data points are colored according to their phylogroup affiliation. The ellipses correspond to 95% confidence intervals for each phylogroup, illustrating the clustering and separation of codon usage profiles among the groups. This analysis highlights the genomic codon usage divergence that correlates with phylogenetic differentiation.

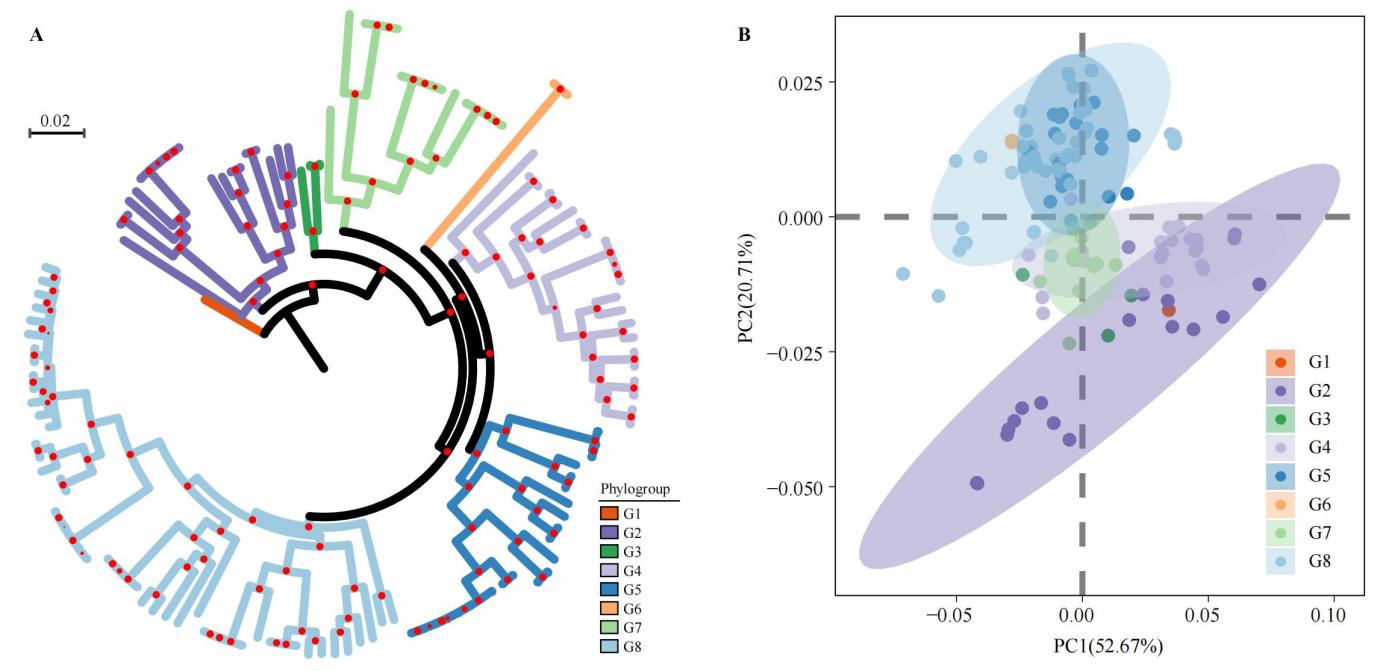

**Figure S6.** Phylogenetic network among phylogroups. The network was constructed using SplitsTree4 based on concatenated nucleotide sequences of core single-copy genes from all analyzed genomes. Branch lengths correspond to genetic distances, with the scale bar representing 0.01 substitutions per site. Different colored shaded regions highlight distinct phylogroups (G1 to G8) to emphasize their clustering within the network. The network structure reveals potential reticulate evolutionary events, such as recombination or horizontal gene transfer, indicated by the presence of network-like connections (splits) rather than a strictly bifurcating tree. This representation complements traditional phylogenetic trees by capturing conflicting signals and highlighting complex evolutionary relationships among the phylogroups.

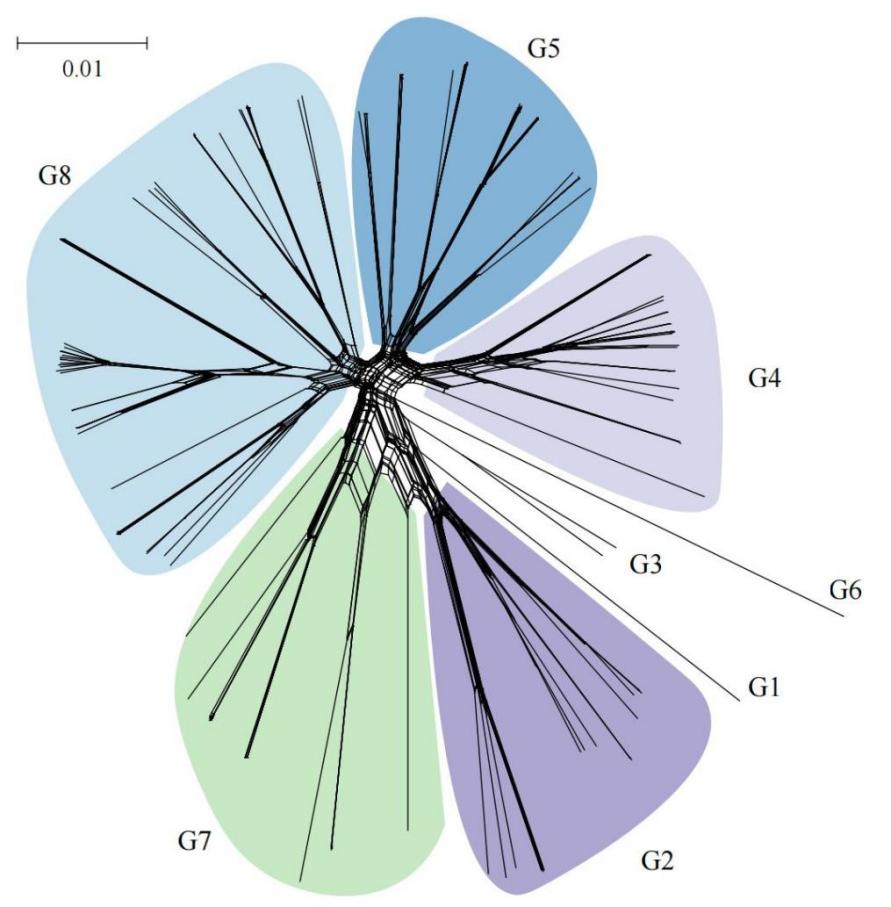

**Figure S7.** Phylogenetic tree based on 16S rRNA gene (*rrs*) sequences. For each strain containing multiple ribosomal operons, the main *rrs* sequence was selected based on the highest sequence similarity and the most significant number of hits compared with sequences from other strains to ensure representativeness. These selected sequences were aligned using a multiple sequence alignment method, and a phylogenetic tree was subsequently constructed using the maximum likelihood algorithm. Branch lengths are proportional to the number of nucleotide substitutions per site, with the scale bar representing 0.005 substitutions per site. Different phylogroups (G1 to G8) are highlighted with distinct colors around the tree circumference for visual differentiation. Red circles at branch nodes indicate bootstrap values above a 75% threshold, reflecting strong statistical support for the corresponding clades.

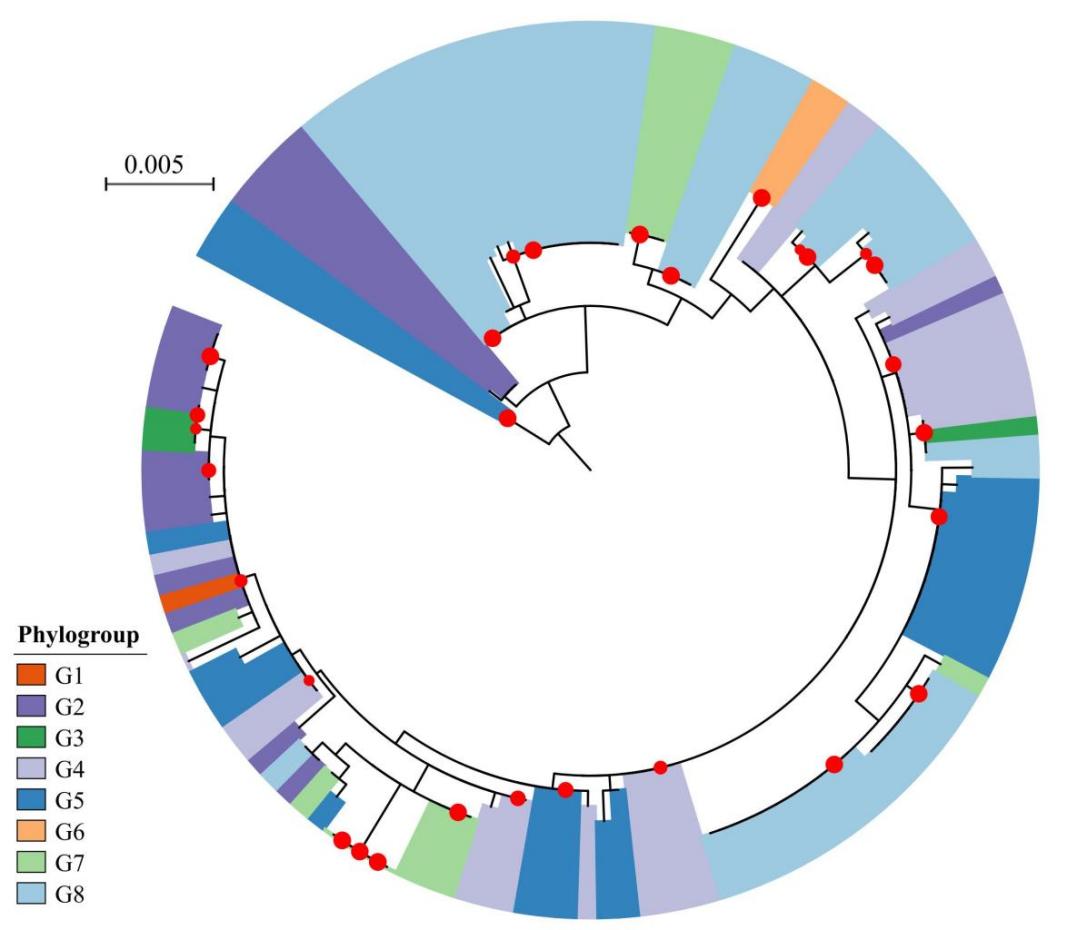

**Figure S8.** Inconsistency of species delimitation based on different thresholds. (A) Heatmap showing pairwise comparisons of 16S rRNA gene (*rrs*) sequence similarity (y-axis) versus average nucleotide identity (ANI) values (x-axis) among the analyzed strains. The heatmap is colored based on commonly used species delineation thresholds: *rrs* similarity greater than 98.5% (indicated in red) and ANI greater than 95% (indicated in red), whereas values below these thresholds are shown in blue. A white diagonal line demarcates comparisons of the same strains. Color blocks along the bottom (ANI) and right side (*rrs*) correspond to phylogroups (G1 to G8), facilitating visualization of clustering consistency between methods. This panel highlights discrepancies where strains meet one threshold but not the other, illustrating challenges in defining species boundaries using different molecular criteria. (B) Plot of the ratio h/m, a metric reflecting gene flow, as a function of increasing genome sample size, starting from smaller datasets (4 genomes) to the complete set including 100 genomes. The outgroup strain *Microcoleus anatoxicus* CHAB4116 is included in the analysis to provide phylogenetic context. The dashed black line represents the mean h/m values, while the shaded gray area indicates the variability around the mean, demonstrating a pronounced mid-range discontinuity and gene flow barrier.

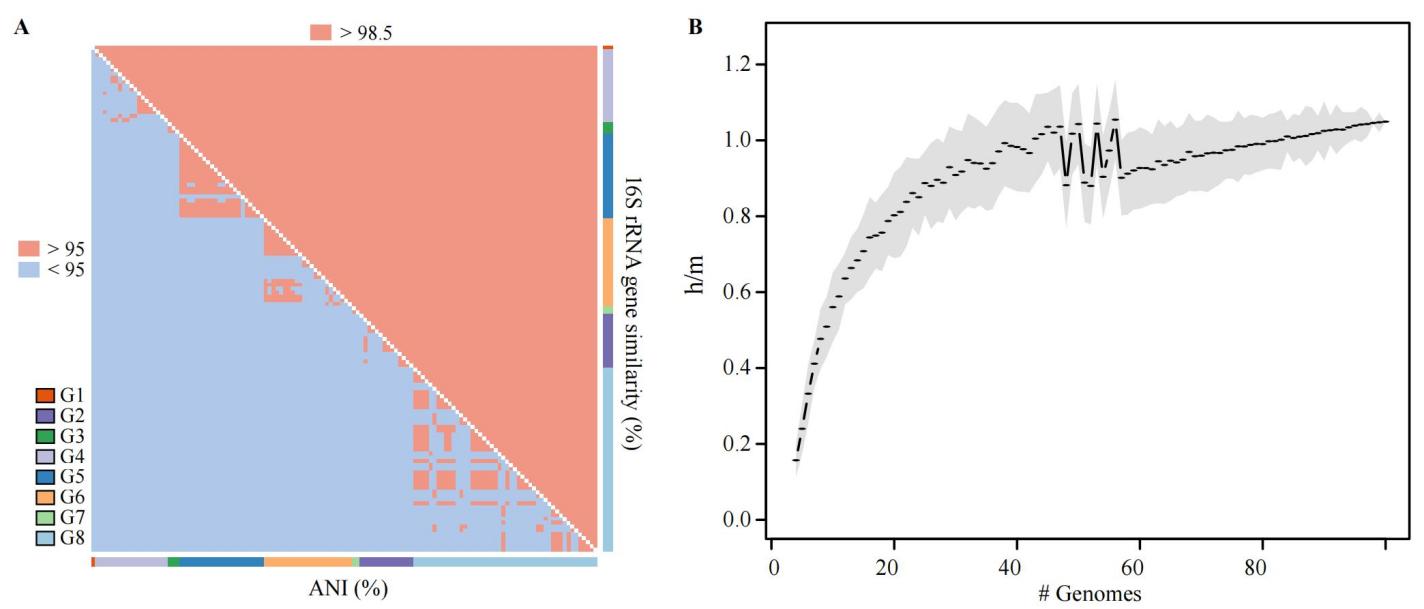

**Figure S9.** Genomic exchange among phylogroups. (A) Sankey diagram illustrating the patterns of ancient and recent gene flow within the core genome across different phylogroups. The flow proceeds from left (ancestral gene flow events) to right (recent gene flow events), with the vertical bars representing phylogroups at each evolutionary timeframe. Colored ribbons depict the direction and extent of gene flow between source and recipient phylogroups, with the thickness of the ribbons proportional to the magnitude of gene exchange. This visualization highlights both historical and ongoing genetic connectivity and differentiation among phylogroups, revealing complex networks of shared genetic material. (B) Circular chord diagram representing the number of inter-phylogroup horizontal gene transfer (HGT) events detected across the entire genome. The outer ring is segmented and color-coded according to specific phylogroups, serving as both donors and recipients. Ribbons connect source and target phylogroups, with the side matching the outer ring color representing the source. The width of each ribbon is scaled to reflect the quantity of HGT events in that direction. This diagram provides a comprehensive overview of recent genetic exchanges, highlighting the asymmetrical dynamics of gene transfer and the preferential pathways among phylogroups within the population.

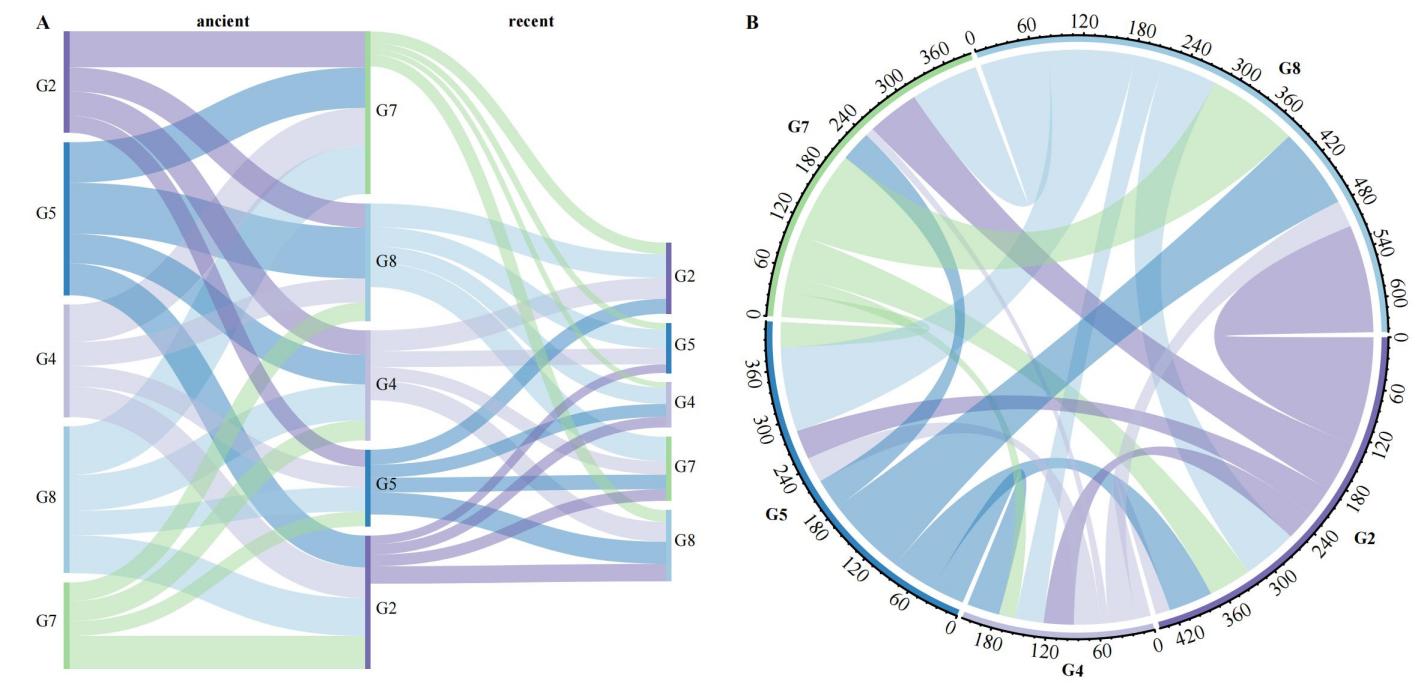

**Figure S10**. Phylogenetic architecture excluding positively selected genes (PSGs). To improve the accuracy and reliability of phylogenetic reconstruction by minimizing the confounding effects of strong adaptive selection, the phylogenetic tree was inferred using protein sequences from core single-copy genes after removal of all identified PSGs. This approach reduces the bias introduced by rapid evolutionary changes in PSGs, which can obscure authentic evolutionary relationships. Branches in the circular cladogram are color-coded according to their respective phylogroups (G1 to G8), as indicated in the accompanying legend, allowing for clear visualization of the clustering patterns and divergence among groups. Black dots on nodes indicate bootstrap support greater than 75%, while branch lengths are proportional to the amount of sequence divergence measured; the scale bar denotes 0.01 substitutions per site.

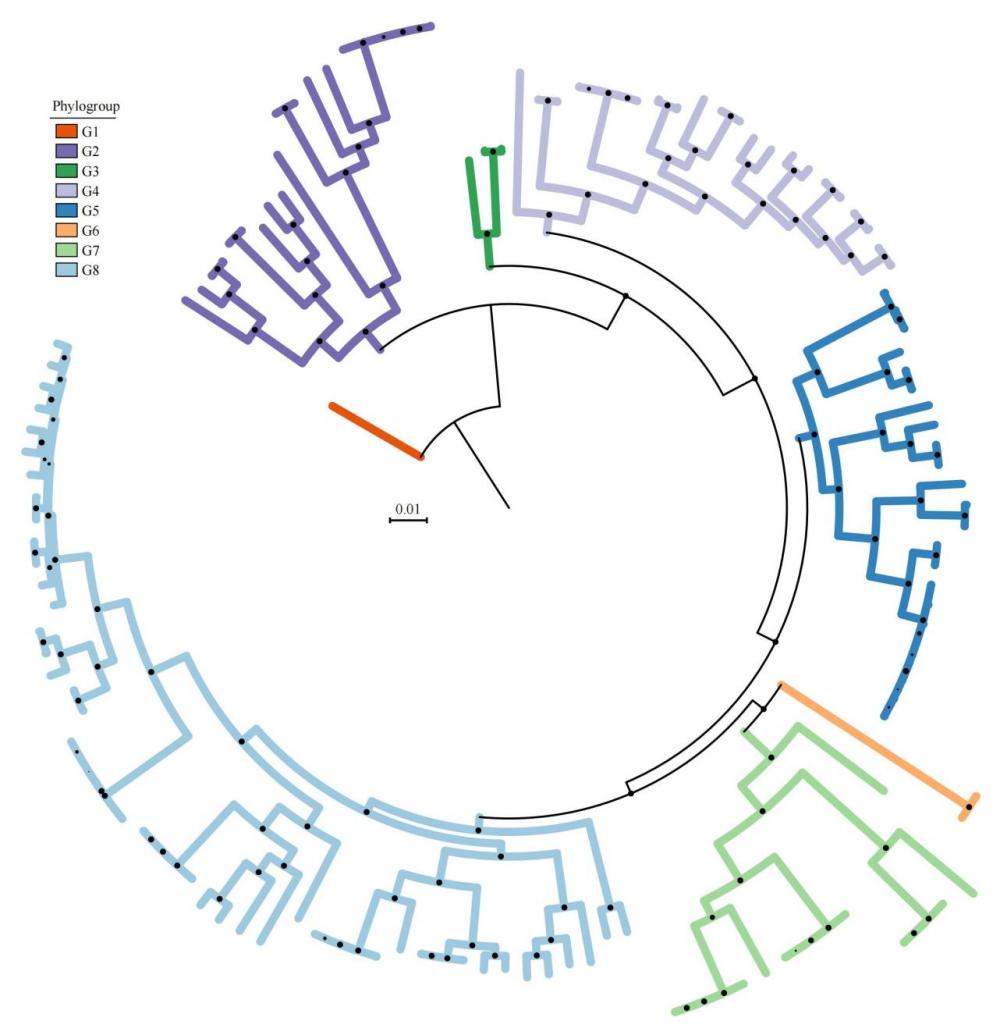

**Figure S11.** Normalized density distribution of nucleotide diversity within genomic regions under positive selection and the total core genome. The figure presents the comparative distributions of nucleotide diversity, measured in 10-kb sliding windows, within the core genome for multiple phylogroups (G2, G4, G5, G7, and G8). Gray-shaded areas depict nucleotide diversity values across all 10 kb windows spanning the entire core genome, reflecting the genome-wide baseline diversity. Orange-shaded areas represent nucleotide diversity within the 10-kb windows containing genes identified as being under positive selection. The comparison across phylogroups reveals divergent evolutionary dynamics associated with selection, highlighting the differential impact of adaptive processes on the genomic architecture among distinct lineages.

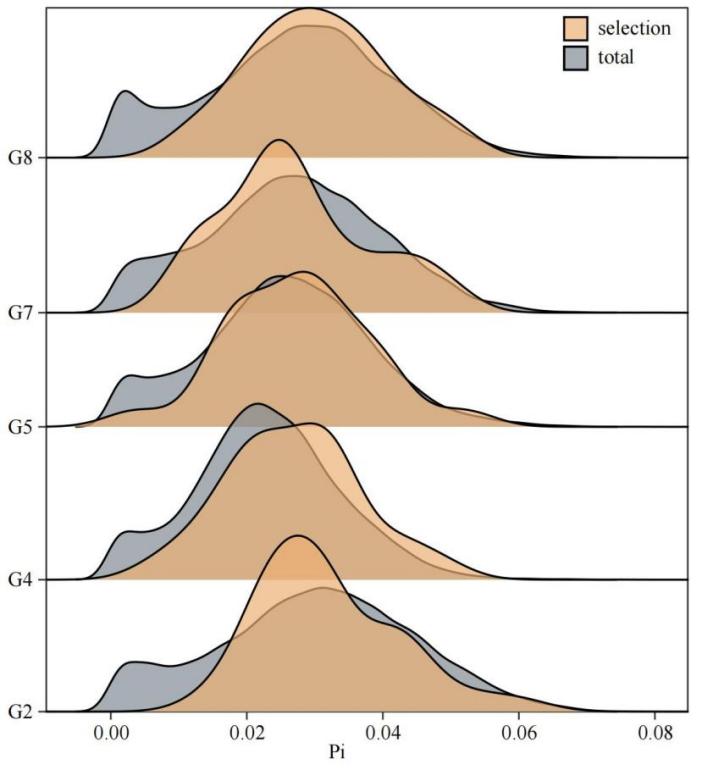

**Figure** **S12**. Relationship between gene family count and phylogenetic distance from the ancestral node across evolutionary nodes within each phylogroup. Each colored line represents a linear regression model fitted to the data points of a specific phylogroup, with shaded areas indicating the 95% confidence intervals. The positive slopes of these lines indicate significant positive correlations between gene family numbers and phylogenetic distance, suggesting gene family expansion along evolutionary divergence. Corresponding p-values and *R*² values for each phylogroup are displayed in the legend, confirming the statistical significance and strength of these correlations. Scatter points represent individual data observations, with slight transparency to reveal data density.

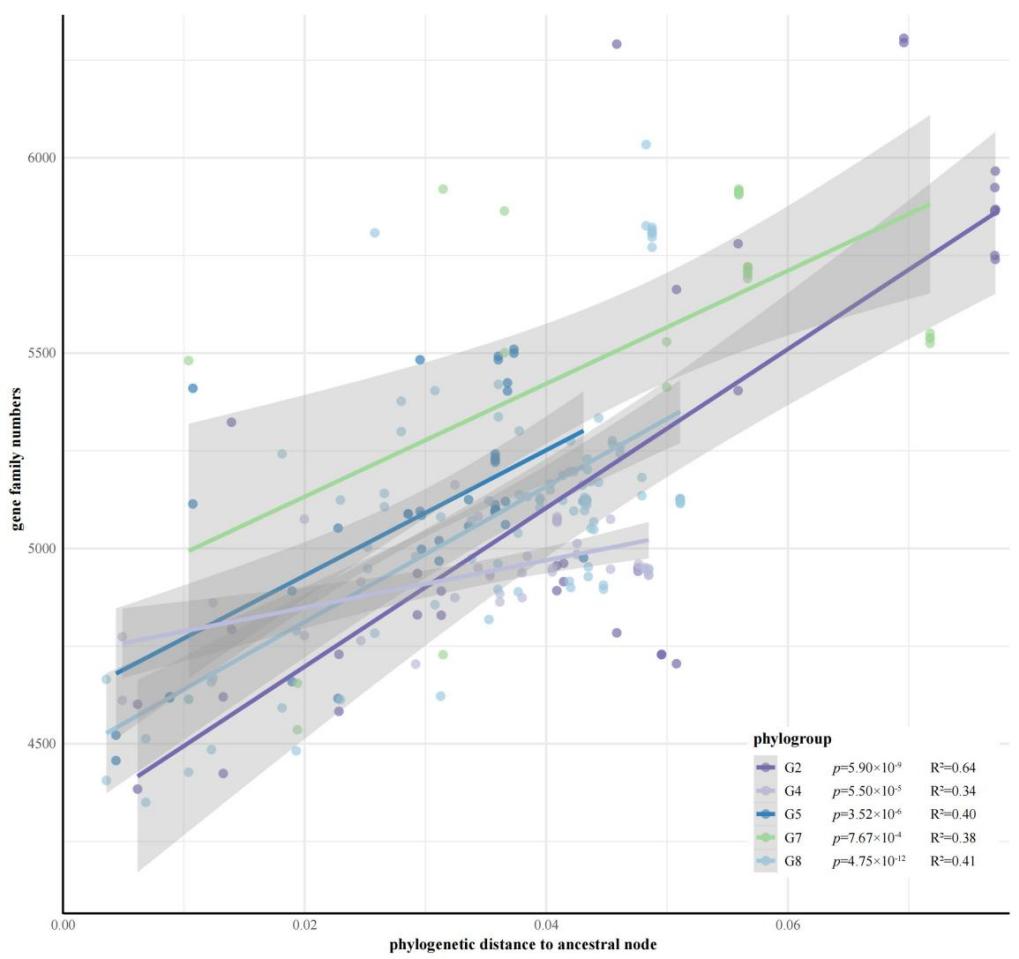

**Figure S13.** Quantile-Quantile (Q-Q) plot of genome-wide association study (GWAS). Each black dot represents a single nucleotide polymorphism (SNP) tested for association with the studied trait. The red diagonal line corresponds to the expected distribution of p-values if there were no true associations, serving as a baseline for comparison. Deviations of points above the diagonal line indicate SNPs with observed p-values smaller than expected, suggesting potential true genetic associations with the trait. This plot effectively visualizes the overall distribution of association signals, helping to distinguish true positives from noise.

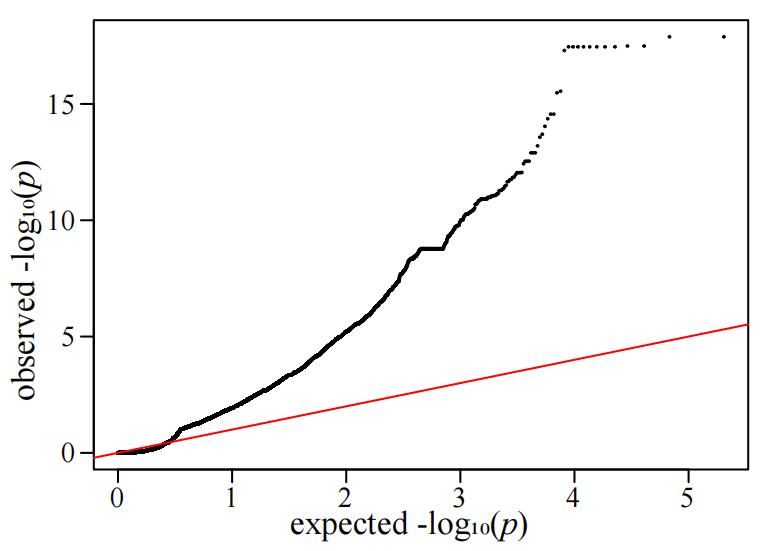

**Table S1.** Comprehensive genomic characteristics and assembly metrics of *Microcoleus vaginatus* strains utilized in this investigation.

**Table S2.** Detailed annotation of genes exhibiting positive selection signatures across phylogroups. Orthologous groups (OGs) marked with asterisks experienced both positive selection and homologous recombination events, while common OGs shared across multiple phylogroups are highlighted with orange backgrounds.

**Table S3.** Comprehensive inventory of genes participating in recombination events across more than one-third of evolutionary nodes within population lineages. An orange background distinguishes common orthologous groups occurring across multiple phylogenetic groups.

**Table S4.** Significantly differentiation-associated SNP loci identified through genome-wide association analysis with the correspondence to previously characterized evolutionary signatures.

**Table S5.** Environmental critical parameter matrix corresponding to individual strains. All values in the matrix were extracted from a public database based on the isolation sites of the strains.
